# Intestinal Barrier Loss Enables Microbiota-Mediated Purinergic Suppression During Malaria

**DOI:** 10.64898/2026.01.22.701153

**Authors:** Rafael B. Polidoro, Olivia J. Bednarski, Sawyer Lehman, Fabrício Marcus Silva Oliveira, Elizabeth M. Fusco, Layne Bower, Marcos Rangel-Ferreira, Sukhmeet Kour, Ruth Namazzi, Robert O. Opoka, Chandy John, Jeffrey J. Brault, Andrew Tilston-Lunel, José Carlos Alves-Filho, Nathan W. Schmidt

**Affiliations:** Ryan White Center for Pediatric Infectious Diseases and Global Health, Herman B. Wells Center for Pediatric Research, Department of Pediatrics, Indiana University School of Medicine, Indianapolis, IN 46202, USA; Department of Microbiology and Immunology, Indiana University School of Medicine, Indianapolis, IN 46202 USA; Department of Microbiology, Immunology and Tropical Medicine, George Washington University, Washington, DC 20052, USA; Laboratory of Innovations in Therapies, Education, and Bioproducts, Instituto Oswaldo Cruz, FIOCRUZ, RJ 21041-250, Brazil; School of Science, Indiana University Indianapolis, IN 46202 USA; Department of Paediatrics and Child Health, Makerere University College of Health Sciences, Kampala, Uganda; Global Health Uganda, Kampala, Uganda; Aga Khan University East Africa Medical College, Nairobi, Kenya; Indiana Center for Musculoskeletal Health, Department of Anatomy, Cell Biology & Physiology, Indiana University School of Medicine, Indianapolis, IN 46202, USA; Department of Pharmacology, Ribeirão Preto Medical School, University of São Paulo, Ribeirão Preto, SP 14049-900, Brazil

**Keywords:** gut microbiota, malaria pathogenesis, regulatory T cells, intestinal barrier integrity, purinergic signaling

## Abstract

The gut microbiota shapes malaria immunity and disease severity, but the mechanisms underlying these effects remain unclear. In a murine model, susceptibility to *Plasmodium yoelii* hyperparasitemia was associated with elevated regulatory T cells and diminished IFN-γ. We demonstrate that the IgA-coated fraction of the microbiota is sufficient to transfer this susceptibility. *Plasmodium* infection disrupts the intestinal barrier regardless of microbiota composition. Mechanistically, barrier loss was associated with systemic adenosine persistence and an expansion of CD39+ plasmablasts in susceptible mice. Ugandan children with severe malaria exhibited a distinct purinergic immune signature compared to asymptomatic community children. Therapeutic reinforcement of the gut barrier or blockade of the downstream adenosine A2A receptor improved germinal centers and reduced disease severity in mice, independent of parasite burden, revealing a purinergic-dependent immunosuppression pathway that drives pathogenesis. This work defines an axis in which malaria-induced gut leakiness enables microbial-derived signals to trigger purinergic immunosuppression and severe disease.

## Introduction

Malaria remains a devastating global health problem, and effective immunity, particularly the generation of long-lived antibody responses, is crucial for protection^1–3^. Systemic immunity against blood-stage *Plasmodium* parasites is a complex process. In recent years, the gut microbiota has emerged as a critical, yet highly variable, environmental factor that can profoundly shape host malaria immunity, gut barrier integrity, secondary bacterial invasion, and parasite burden ^4–9^. Our previous work highlighted this variability, demonstrating that inbred mice from different commercial vendors exhibit starkly different susceptibilities to *P. yoelii* 17XNL hyperparasitemia, a phenotype driven by their distinct gut microbial compositions^10,11^. This observation is not limited to murine models; specific bacterial taxa in African children, such as *Bacteroides*, are associated with an increased risk of severe malaria^12^, and bacterial dysbiosis during severe malaria contributes to its pathogenesis in African children^13^. Moreover, pre-malaria transmission-season gut bacterial composition in Malian children was associated with susceptibility to febrile malaria during the ensuing transmission season^14^. Finally, germ-free mice colonized with stool samples from Malian children susceptible to febrile malaria showed increased parasite burden compared with those colonized with stool samples from children resistant to febrile malaria^14,15^.

A key mechanism through which the gut microbiota exerts this systemic influence is by modulating the quality of the B cell response. We previously established that microbiota-driven susceptibility to *P. yoelii* hyperparasitemia is associated with a defective germinal center (GC) reaction, characterized by premature contraction of GCs, and a more narrow and variable anti-*P. yoelii*-specific antibody repertoire^16,17^. This ultimately leads to a compromised B-cell memory response, hindering long-term protection^16,18^. However, the precise cellular and molecular pathways by which specific gut microbial communities disrupt this critical GC process have remained unclear.

In this study, we sought to dissect the mechanistic links between the gut, host immunity, and malaria pathogenesis. We hypothesized that a “susceptible” microbiota establishes a tolerogenic immune environment, marked by an altered regulatory T cell (Treg) response, that becomes pathogenic upon the onset of *P. yoelii* infection. We show that susceptibility to hyperparasitemia is associated with a failure to downregulate splenic and intestinal Tregs during infection and a blunted systemic cytokine response. Although infection induces intestinal barrier disruption in both resistant and susceptible mice, the pathological outcome might be dictated by the specific microbial community. Indeed, we find that the IgA-coated fraction of the microbiota from susceptible mice is sufficient to transfer the disease phenotype to resistant animals. This pathology culminates in a shift towards an immunosuppressive, extrafollicular B cell response characterized by high expression of the immunoregulatory ectonucleotidase CD39. This immune-purinergic signature is also observed in Ugandan children with severe malaria. Finally, we demonstrate that disease severity can be uncoupled from parasite burden in mice by therapeutically targeting intestinal barrier integrity or the purinergic A2A receptor, restoring the GC response and improving survival.

## Results

### A microbiota-driven Treg response is linked to malaria susceptibility and a blunted cytokine profile

To define the mechanisms underlying gut microbiota-dependent susceptibility to malaria, we used our established murine model, comparing hyperparasitemia-resistant “resistant” and susceptible “susceptible” C57BL/6 mice. While there were no significant differences in parasitemia between male and female mice, allowing for the use of female mice in subsequent experiments (**Fig. 1A**), susceptible mice exhibited significantly higher parasite burdens and approximately 25% mortality. In contrast, all resistant mice survived the infection (**Fig. 1B**).

**Figure 1.**
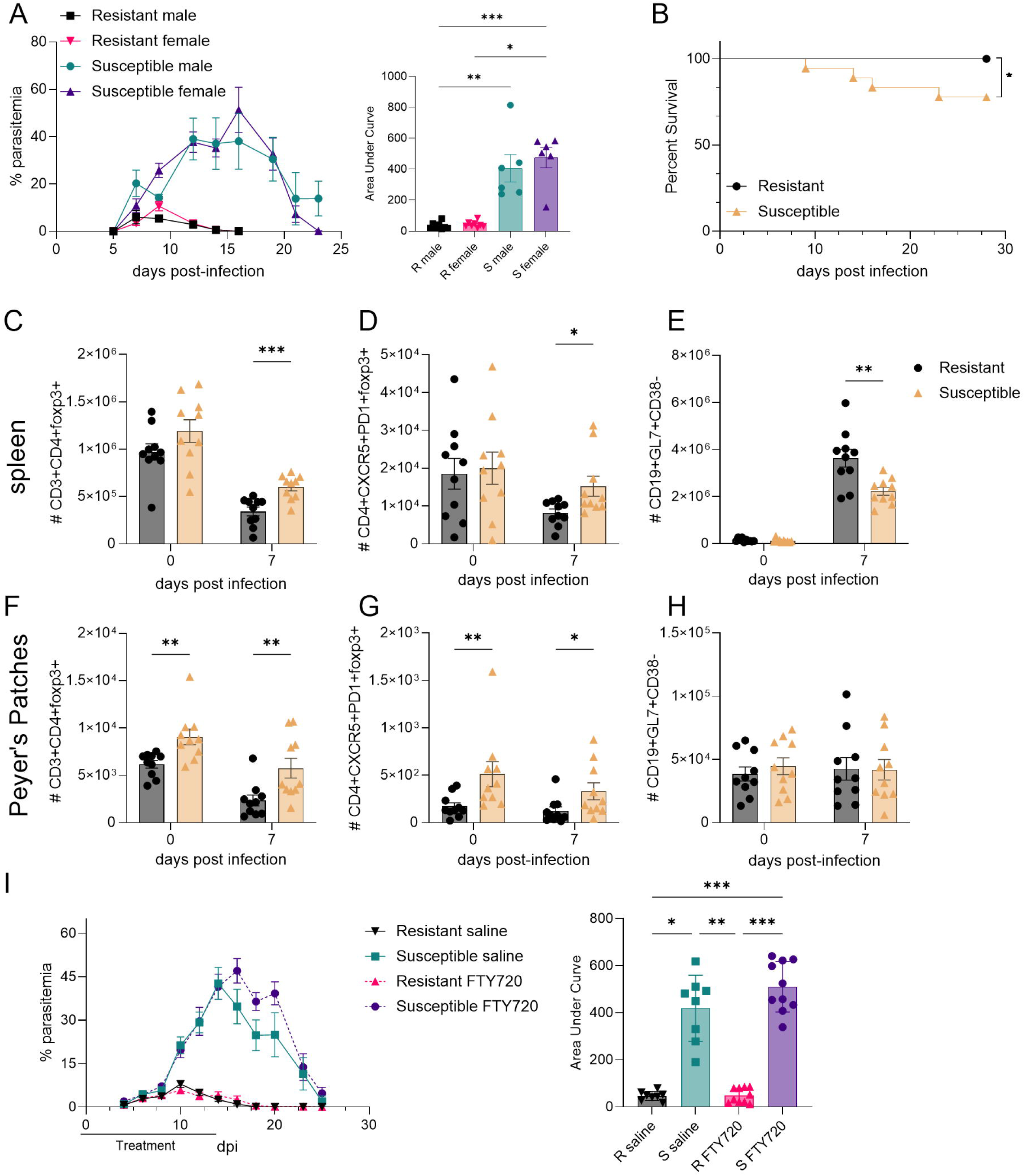
Susceptibility to hyperparasitemia is associated with a systemic regulatory T cell phenotype. (**A**) Longitudinal blood parasitemia (left) and Area Under Curve (right) of C57BL/6 mice (N=6-8) infected with *P. yoelii*. (**B**) Survival curves compared using the Mantel-Cox Log-Rank test. (**C**-**E**) Spleens from Resistant and Susceptible mice were processed for flow cytometry before infection (0) and 7 days p.i. Quantification of (**C**) Tregs (CD3+CD4+Foxp3+Non-Tfr), (**D**) Tfr cells (CD4+CXCR5+PD1+Foxp3+), and (**E**) GC B cells (CD19+GL7+CD38-). (**F-H**) Peyer’s Patches quantification of (**F**) Tregs, (**G**) Tfr cells, and **(H)** GC B cells. (**I**) Parasitemia (left) and Area Under Curve (right) of Resistant and Susceptible mice treated with FTY720 (1 mg/kg) or saline every other day from d-1 to d14. Statistics: Mann-Whitney tests. N=10, Mean ± SEM. *p<0.05, **p<0.01, ***p<0.001, ****p<0.0001

Given our previous work linking susceptibility to impaired germinal center (GC) responses, we hypothesized that regulatory T cells (Tregs) and T follicular regulatory (Tfr) cells might be dysregulated. We analyzed immune cell populations in the spleen, Peyer’s Patches (PPs), and mesenteric lymph nodes (MLNs) by flow cytometry before infection (day 0) and at day 7 post-infection (dpi), a time point immediately preceding the GC contraction we previously observed in susceptible mice^16^ (representative gating in **Suppl. Fig. 1B**). While splenic Treg (CD4⁺Foxp3⁺CXCR5^-^PD1^-^) and Tfr (CD4⁺CXCR5⁺PD1⁺Foxp3⁺) cell counts were comparable between resistant and susceptible mice before infection, they were significantly elevated in susceptible mice at 7 dpi compared to resistant mice (**Fig. 1C**, **D**). This increase in regulatory cells coincided with a significant reduction in GC B cells (CD19⁺GL7⁺CD38⁻) in the spleens of susceptible mice at the same time point (**Fig. 1E**).

To determine if this regulatory phenotype was localized to the spleen or reflected a more systemic state, we examined gut-associated lymphoid tissues. We observed a similar pattern of elevated Tregs in the PPs and MLNs of susceptible mice (**Fig. 1F**, **G**; **Suppl. Fig. 1C**). However, consistent with *P. yoelii* not being a primary intestinal infection, we observed no significant changes in the GC B cell response in the PPs of either group (**Fig. 1H**). The elevated Treg presence in the spleen and gut-associated lymphoid tissue suggested that continuous trafficking of T cells from the gut might suppress the splenic response. To test this hypothesis, we treated mice with FTY720 (fingolimod), a compound that blocks lymphocyte egress from secondary lymphoid organs^19^. However, this treatment did not alter the course of parasitemia in either the resistant or susceptible groups (**Fig. 1I**), suggesting the divergent immune phenotypes are not maintained by continuous T cell migration during an established infection.

Building on the finding of an expanded Treg population, we next evaluated the cytokine environment that shapes the adaptive immune response. We measured key cytokines in spleen homogenates and serum at 5 and 7 dpi. Resistant mice mounted a more robust IFN-γ response, with significantly higher levels in both spleen supernatants and serum compared to susceptible mice (**Suppl. Fig. 1D**). Similarly, IL-10 and IL-4, both of which support GC reactions, were also significantly elevated in the spleens of resistant mice (**Suppl. Fig. 1E-F**)^20,21^. In contrast, levels of the critical T follicular helper (Tfh) cell cytokine, IL-21, were not different between the groups (**Suppl. Fig. 1G**).

Together, these data demonstrate that susceptibility is associated with a systemic expansion of Tregs and a concomitant failure to produce cytokines required to sustain GCs. This suppressive environment appears to be driven by signals distinct from active cell trafficking, as blocking lymphocyte egress did not affect disease outcome. We therefore hypothesized that the mechanism connecting the gut microbiota to this splenic suppression involves a breakdown of the intestinal barrier, allowing microbial signals to leak systemically.

### *P. yoelii* infection induces intestinal pathology and tight junction disruption irrespective of susceptibility phenotype

The distinct basal Treg populations in the gut of resistant and susceptible mice suggested a functionally different intestinal immune environment. To test this, we challenged uninfected mice with a sub-colitis dose of 2% dextran sulfate sodium (DSS), a widely used chemical agent that disrupts the epithelial barrier and induces inflammation. As hypothesized, mice susceptible to hyperparasitemia, which harbor higher intestinal Treg levels, were significantly more resistant to DSS-induced weight loss than hyperparasitemia-resistant Tac mice (**Fig. 2A**). This confirmed that distinct microbiota compositions confer opposing susceptibilities to distinct inflammatory challenges. Importantly, the transient dysbiosis caused by DSS treatment did not durably alter the malaria phenotype, as recovered mice showed no changes in parasitemia, weight loss, or mortality upon subsequent *P. yoelii* infection compared to mice not previously treated with DSS (**Suppl. Fig. 2A**-**C**).

**Figure 2.**
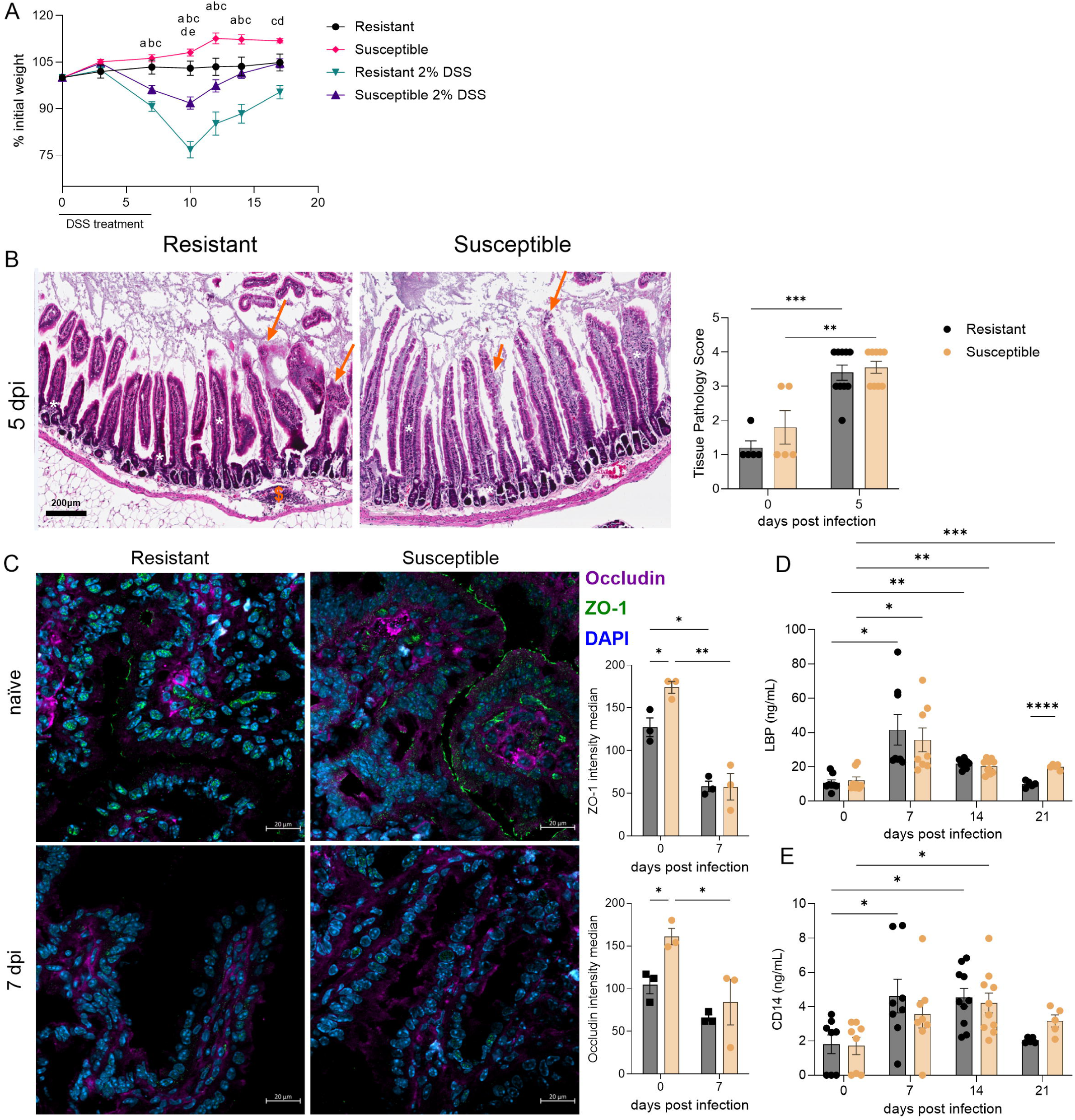
Gut microbiota confers resistance to DSS colitis but does not prevent infection-induced barrier damage. (**A**) Resistant and Susceptible mice were treated with 2% DSS for 7 days. Body weight percentage relative to initial weight. (**B**) Representative H&E staining of small intestine sections at day 5 post-*P. yoelii* infection showing mucosal desquamation (arrow), inflammatory infiltrate in the mucosa (*) and submucosa ($) (scale bar: 200µm) and pathology scores (right). (**C**) Immunofluorescence of the distal small intestine stained for Occludin (purple), ZO-1 (green), and DAPI (blue). White bar 20 µm. Right: Median fluorescence of 40x objective images of intestine or region of interest when the lumen was abundant was measured using Zen Blue Lite. Mean ± SEM. (**D-E**) Longitudinal serum quantification of **(D)** LBP and **(E)** sCD14 by ELISA on days 0, 7, 14, and 21 p.i. Statistics: Mann-Whitney (pathology/IFA) and One-way ANOVA/Mixed-effects (longitudinal). Fig 2A comparisons p>0.05: a – Resistant vs. Resistant 2% DSS; b - Resistant 2% DSS vs. Susceptible; c – Susceptible to Susceptible 2% DSS; d – Resistant 2% DSS to Susceptible 2% DSS; e – Resistant to Susceptible 2% DSS. *p<0.05, **p<0.01, ***p<0.001. DSS data are representative of 4 experiments. Pathology scores and sections are accumulated from two independent experiments.

This led us to question whether the severity of intestinal damage during malaria infection itself differed between the groups. We performed histopathological analysis of the small intestine before and after infection. While naive animals showed normal histology, resistant and susceptible mice developed comparable, moderate-to-intense inflammatory infiltrates in the mucosa and submucosa by 5 dpi (**Fig. 2B**, **Suppl. Fig. 2D**). To investigate the basis of this pathology, we examined the integrity of the intestinal barrier. Immunofluorescence analysis revealed a dramatic, infection-induced disruption of the key tight junction proteins ZO-1 and occludin in the distal small intestine of both groups at 7 dpi (**Fig. 2C**, **Suppl. Fig. 2E**). To confirm that this local intestinal damage led to systemic consequences, we measured serum biomarkers and found that markers of bacterial translocation sCD14 and LBP (**Fig. 2D-E**), and vascular damage Angiopoietin-2 and ICAM-1 (**Suppl. Fig. 2F**) were all significantly elevated in both resistant and susceptible mice following infection. These results indicate that *P. yoelii* infection is associated with intestinal inflammation, tight junction disorganization, and increased systemic biomarkers regardless of host predisposition to hyperparasitemia.

### The IgA-coated fraction of the microbiota is sufficient to transfer susceptibility

To test the functional relevance of the immunologically active microbial community^22^. We performed IgA+ enrichment of the cecal contents of susceptible animals and transplanted them into hyperparasitemia-resistant mice. Strikingly, IgA-coated bacteria from susceptible donors were sufficient to render them susceptible to *P. yoelii* hyperparasitemia and to suppress splenic GC responses (**Fig. 3A-D**). This effect was specific, as the IgA-negative flow-through (FT) fraction did not significantly alter the disease course, and IgA-coated bacteria from resistant donors failed to transfer susceptibility (**Suppl. Fig. 3A**).

**Figure 3.**
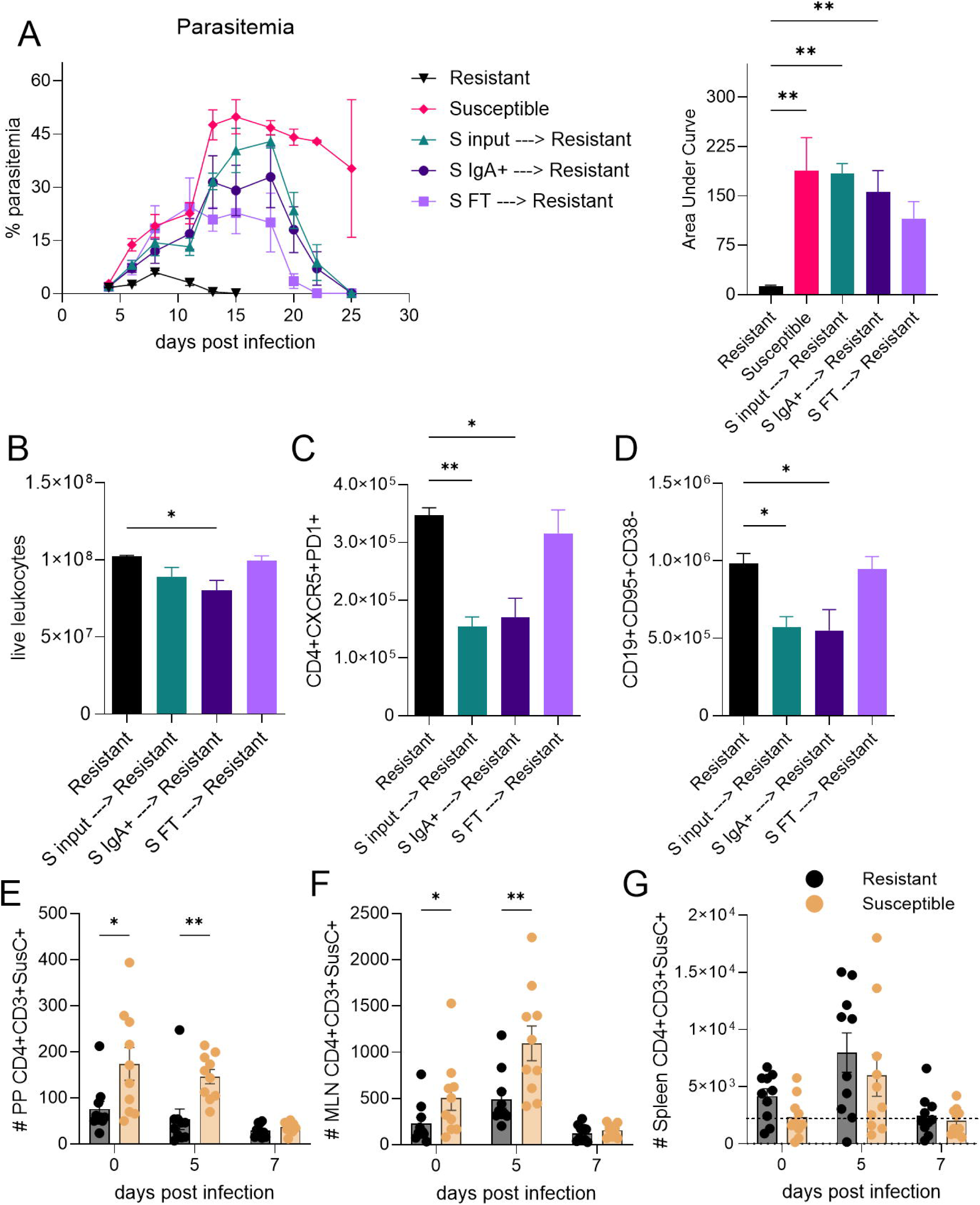
Transfer of Susceptible-IgA+ bacteria into resistant mice is sufficient to cause susceptibility to hyperparasitemia. Cecal contents from naïve susceptible mice were incubated with anti-IgA antibodies and subsequently captured using magnetic beads. Mouse Anti-IgA-PE-bound product (S IgA+), the flow-through (S FT) of the enrichment process, or input content from Susceptible mice ceca were gavage into Resistant mice during three consecutive days a week before infection with *P. yoelii*. Negative Resistant control received saline, and Susceptible mice were used as littermate controls. (**A**) Parasitemia was assessed for all groups, and the area under the curve was calculated. Twenty-three days post-infection, mice splenocytes were subjected to immunophenotyping of live single leukocytes (**B**), T follicular CD4 T cells (**C**), and GC B cells (**D**). Cells from the Peyer Patches (**E**), mesenteric lymph node (**F**), and splenocytes (**G**) of infected and uninfected mice were incubated with tetramers loaded with *B. theta* SusC-like or scramble peptides and subjected to flow cytometry. The data are representative of two independent experiments. Mean ± SEM. IgA-gavage experiments, Ordinary One-Way ANOVA compared to Resistant saline controls (**A**-**D**). Tetramer (**E**-**G**) Mann-Whitney comparison. *p<0.05, **p<0.01

Having established that the IgA-coated fraction was sufficient to transfer susceptibility, we next sought to identify the specific taxonomic composition of the immunomodulatory consortium. We performed whole-genome sequencing of the cecal microbiota from susceptible mice, separating the community into IgA-coated and non-coated fractions. The total microbial community (input) was dominated by *Bacteroides thetaiotaomicron* (**Suppl. Fig. 3B**), a commensal known to have anti-inflammatory properties in intestinal disease models, particularly impacting CD4+ T cells^23,24^. Consistent with an active, local immune response to this abundant species, we observed elevated numbers of *B. thetaiotaomicron*-reactive T cells (SusC-like tetramer+)^23^ in the Peyer’s Patches and MLNs of susceptible mice, but not in the spleen (**Fig. 3E**-**G**). The IgA-coated fraction was enriched for immunomodulatory species such as *Lactobacillus murinus*, used in probiotics, *Candidatus arthromitus*, and members of the *Clostridiaceae* and *Lachnospiraceae* families, rather than overt pathogens^25,26^ (**Suppl. Fig. 3B**). These findings suggest that the suppression of host immunity is a broader characteristic of the microbiota, not a trait limited to specific pathogens.

### Pharmacological blockade of intestinal permeability ameliorates severe malaria and restores protective immunity

Our findings suggest that infection-induced intestinal barrier dysfunction is a critical event that enables detrimental gut-spleen crosstalk. We therefore treated susceptible mice with larazotide acetate (LA), a zonulin antagonist that reinforces tight junctions^27,28^, after the onset of intestinal damage (days 6-10 post-infection). LA treatment significantly reduced intestinal permeability in infected mice, as measured by plasma levels of 4 kDa FITC-dextran (**Fig. 4A**).

**Figure 4.**
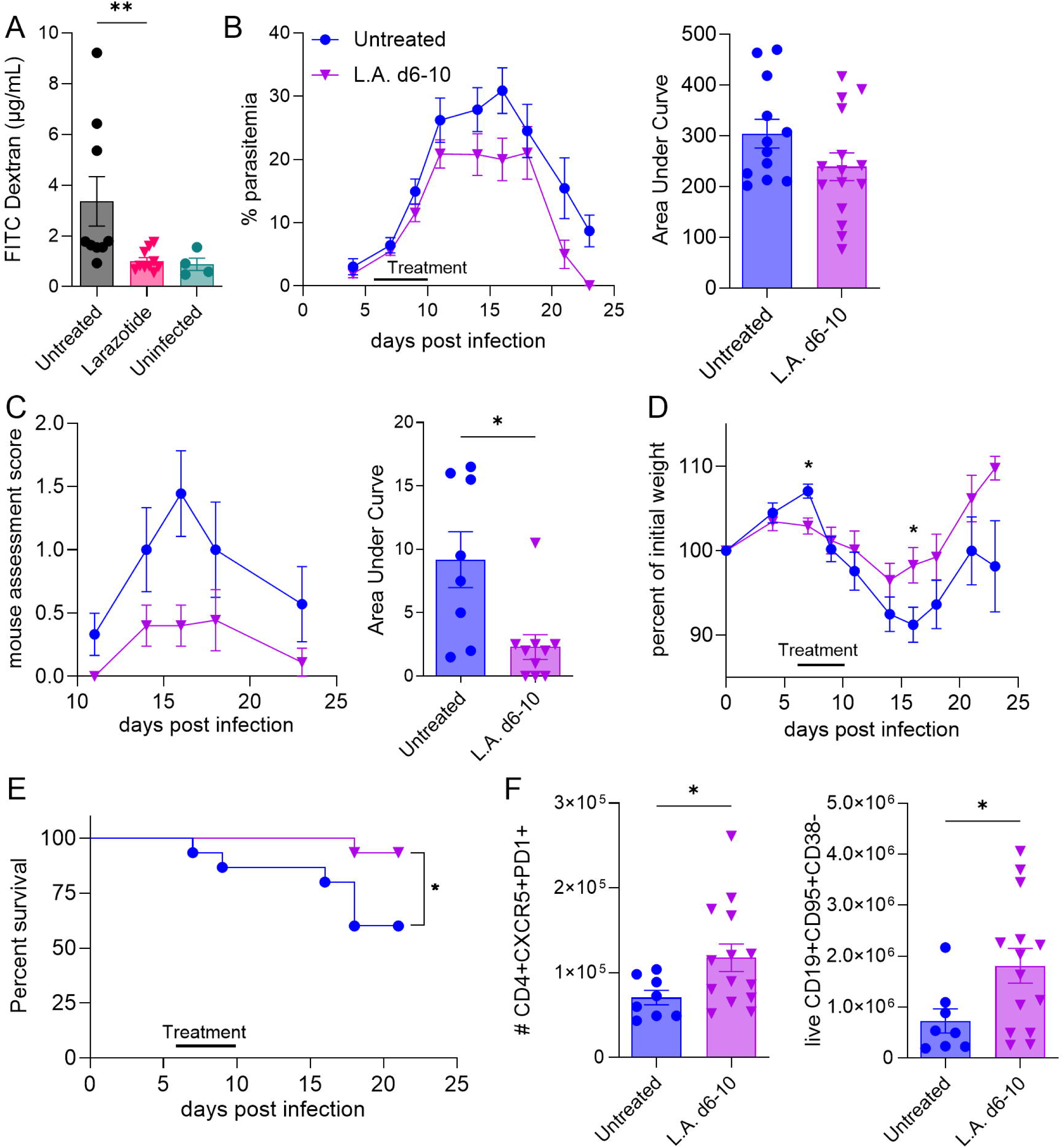
Antagonizing the gut permeability regulator zonulin improves severe malaria symptoms in mice. 8-week-old C57BL/6 mice were infected with *P. yoelii* and treated with L.A. 0.15 mg/mL by gavage (**A**) and/or in drinking water (**A-F**), from d6-d10 post-infection. (**A**) Mice were starved for 4 hours, gavaged with FITC-Dextran 4 kDa (F4k), and plasma was obtained, diluted 1:1 in PBS, and compared to the normal mouse serum F4k dilution curve. (**B**) Parasitemia, d5-16 area under the curve of parasitemia, and survival. (**C**) Sickness scores and areas under the curve of the scores: An independent researcher blindly administered the sickness scores and measured activity levels for state, dehydration, hunching, closing eyes, and the capacity to return to balance. (**D**) % of initial weight assessments. (**E**) Survival curves were compared using the Mantel-Cox Log-Rank test. (**F**) Spleens were collected at 23 dpi. Cells were gated as single cells, then as lymphocytes, then as live (Zombie Aqua negative) CD3+, and then gated as depicted for Tfh CD4 T cells and GC B cells. Pairwise comparisons for area under the curve and cell frequencies were compared using the Mann-Whitney unpaired test (two-tailed). Initial N=15 except for **A** and **C** (N=10 for untreated and L.A., N=4 for uninfected).

While LA treatment only had a modest, non-significant effect on parasite burden (**Fig. 4B**), it dramatically impacted host health, significantly reducing sickness scores, protecting from severe weight loss, and increasing survival from 66% to over 93% (**Fig. 4C**-**E**, **Suppl. Fig. 4A**). This clinical recovery was associated with a mild restoration of the adaptive immune response; spleens from LA-treated mice on day 23 post-infection contained significantly more Tfh and GC B cells compared to controls (**Fig. 4F**). The therapeutic window for this intervention was critical, as treatment initiated earlier (days 1-10 or 4-10) failed to confer protection, likely due to compensatory mechanisms outcompeting the drug during the onset of intestinal damage (**Suppl. Fig. 4**). These data demonstrate that pharmacologically maintaining intestinal barrier integrity during acute infection can prevent severe disease and improve a late anti-malarial GC response.

### Susceptibility is marked by a shift to an emergency B cell response with high CD39 expression

Biomarkers of microbial product translocation following intestinal barrier disruption suggest a sepsis-like pathology may contribute to the observed immunosuppression. Given the role of purinergic signaling in sepsis^29^, we investigated the expression of the ectonucleotidase CD39 in our model. The purinergic system is present in a balance between pro-inflammatory effector functions and anti-inflammatory resolution^30^. A central mechanism is the CD39/CD73-dependent conversion of pro-inflammatory extracellular ATP into immunosuppressive adenosine^31,32^. Unbiased FlowSOM^33^ metaclustering of B220⁺ B cells at 7 dpi revealed a stark divergence in the B cell response: susceptible mice displayed a massive expansion of CD138^+^IgM^+^ B cells consistent with plasmablasts (**Fig. 5A**-**C**). Critically, CD138^+^ populations 5 and 6 exhibit markedly higher CD39 expression independent of the susceptibility to hyperparasitemia (**Fig. 5C**, dot plot heatmap). We confirmed these findings using conventional flow cytometry gating (gating strategy in **Suppl. Fig. 5A**). Susceptible mice had significantly more IgM⁺ and IgM⁻ plasmablasts (CD19^+^CD138^+^B220^lo^) at 7 dpi (**Fig. 5D**-**E**). Following infection, the CD39 MFI was significantly higher in susceptible mice than their resistant counterparts (**Fig. 5F**, **G**). A similar pattern was observed in Tregs (**Suppl. Fig. 5B**).

**Figure 5.**
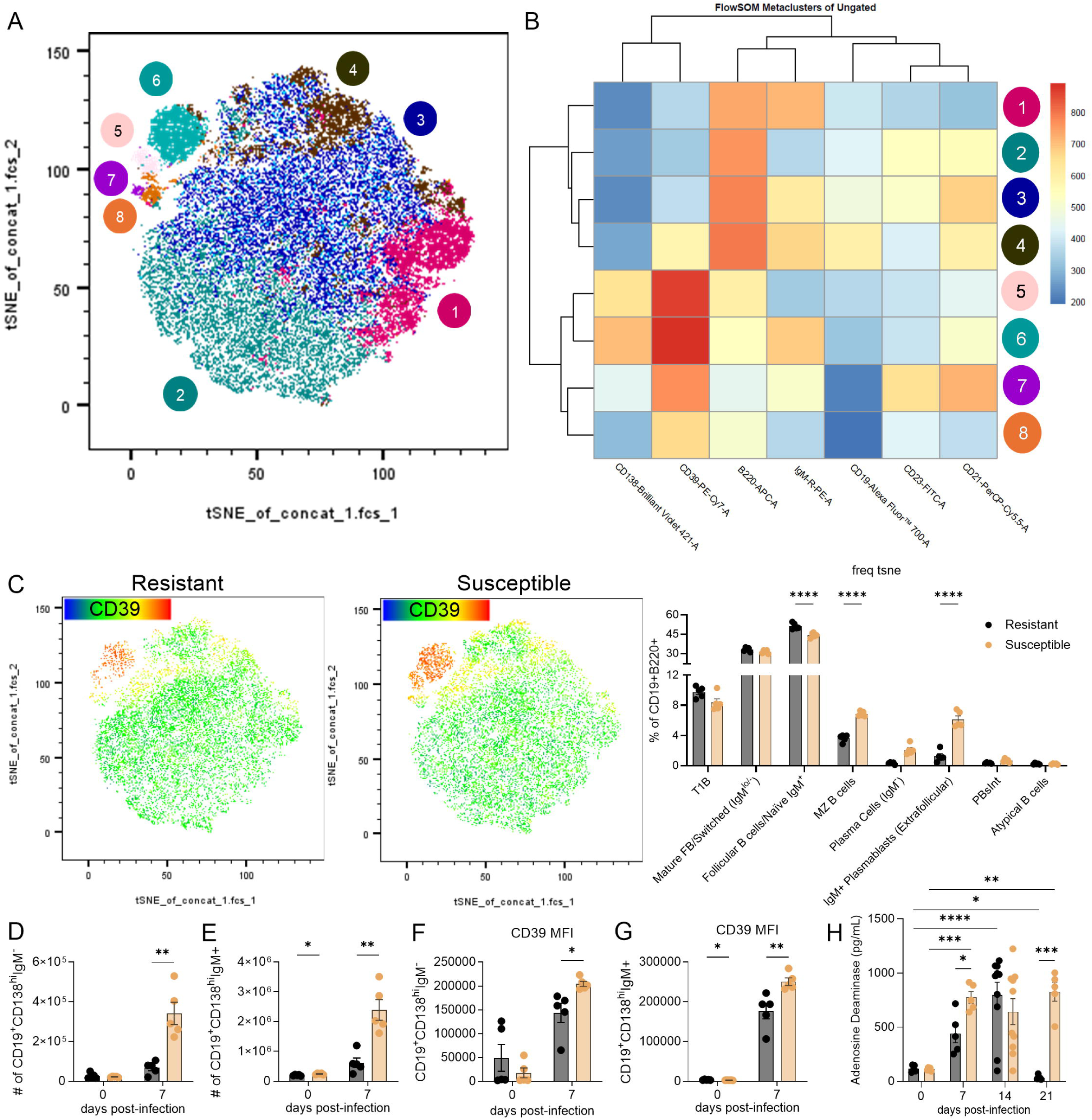
Susceptibility is characterized by high CD39 expression on plasmablasts and elevated systemic ADA levels. Spleen from 5 Resistant and Susceptible uninfected and 7-day *P. yoelii*-infected mice were harvested, processed, and subjected to flow cytometry after staining for CD39 and B and Treg cell markers. (**A**) Unsupervised FlowSOM clustering in B-cell-stained panel applied over t-SNE, showing eight distinct populations for 10,000 singlet live CD19+ cells for each mouse. (**B**) Unsupervised Heatmap Metaclusters of FlowSOM of the populations. (**C**) CD39 Heatmap representation of Resistant and Susceptible mice over t-SNE, and the bar graph representing the individual frequencies of the populations with rough annotations based on their heatmap expression level in numerical order. (**D**-**G**) Live singlets were gated as depicted in the Y axis in CD3^-^CD19^+^ Plasmablasts IgM-(**D**), Plasmablasts IgM+ (**E**), and the median fluorescence intensity (MFI) for the CD39 receptor in the same populations (**F**-**G**). (**H**) Longitudinal serum adenosine deaminase (ADA) activity (pg/mL) in Resistant and Susceptible mice. The data are representative of 2 experiments. *p<0.05, **p<0.01, ***p<0.001, ****p<0.0001. Median ± SEM.

To determine whether the host counteracts this potential adenosine accumulation derived from CD39 activity, we measured adenosine deaminase (ADA) levels, the enzyme responsible for degrading adenosine^34^. We observed that susceptible mice mounted a significantly higher serum ADA response at 7 dpi compared to resistant mice (**Fig. 5H**). Furthermore, while ADA levels returned to baseline in resistant mice by 21 dpi, they remained elevated in susceptible animals, consistent with prolonged parasitemia. These results indicate that *Plasmodium* infection activates the purinergic axis, a response that is significantly exacerbated in the context of microbiota-dependent susceptibility.

### Severe malaria induces a purinergic-dependent, emergency immune response in humans

To determine if the purinergic axis and immune paralysis we observed in mice is relevant in human disease, we analyzed single-cell CITE-seq data from a cohort of Ugandan children, previously described^35^, with severe malaria (SM; n = 8) compared to community children (CC; n = 8), half of which had asymptomatic *Plasmodium falciparum* infection, using the Seurat pipeline^36^. There were significant differences across many activated immune populations, with monocytes (CD14 or CD16 annotated) showing the greatest difference in children with SM (**Fig. 6A**, **Suppl. Fig. 6A**).

**Figure 6.**
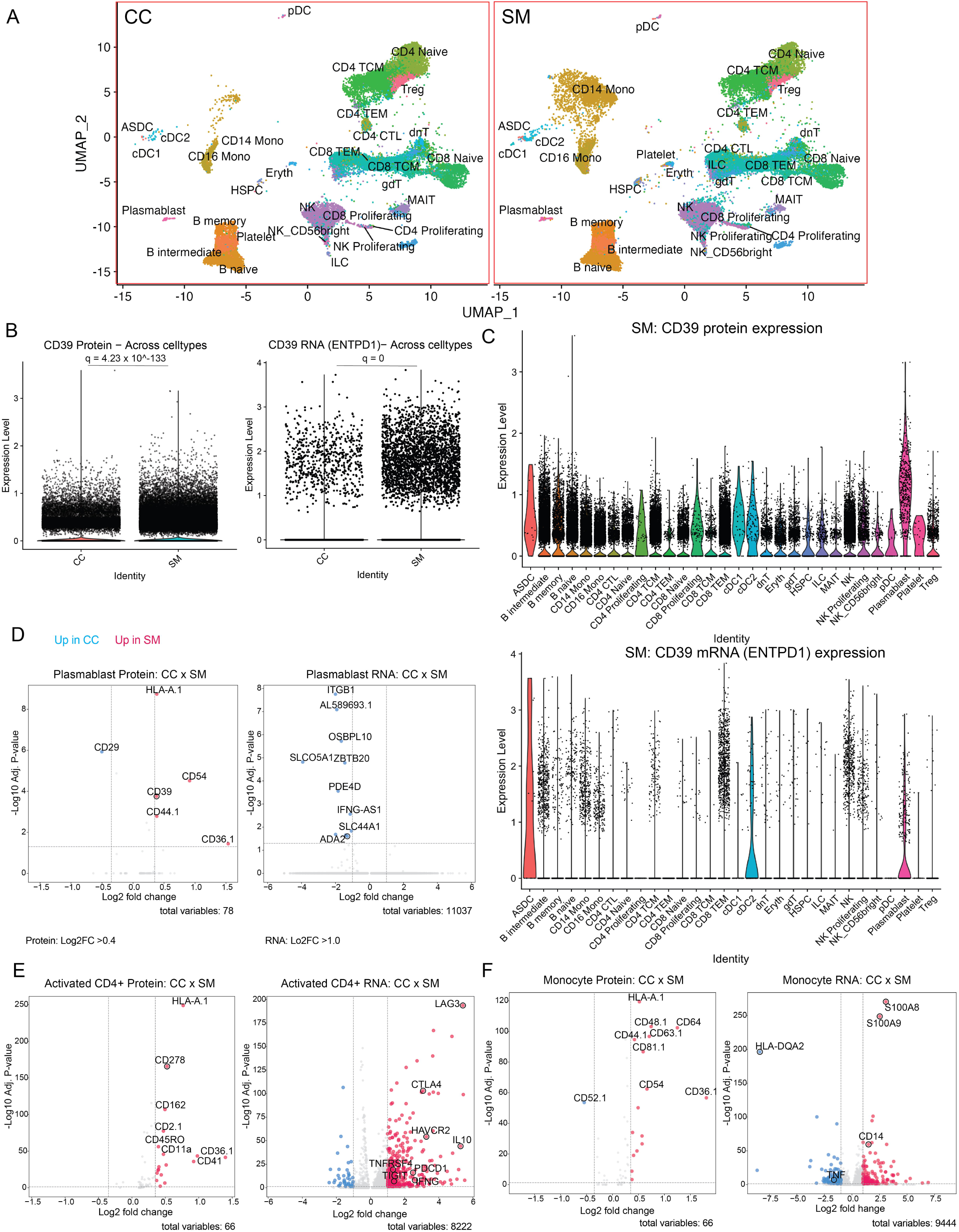
Severe malaria induces an emergency, purinergic-dependent immune response in humans. (**A**) Split UMAP visualization of human PBMCs from healthy community children (CC, left) and children with severe malaria (SM, right). (**B-C**) Plots quantifying normalized CD39 protein or mRNA (ENTPD1) expression across combined (**B**) or individually annotated (**C**) cell types in SM/CC. (**D**-**F**) Volcano plots representing differentially expressed surface proteins (left columns) and mRNA transcripts (right columns) for (**D**) Plasmablasts, (**E**) Activated CD4+ T cells (selected by LAG3^+^CD45RO^+^), and (**F**) Monocytes comparing Severe Malaria (SM) versus Community Children (CC). Red and blue dots indicate significantly upregulated features in SM and CC, respectively. Significance was defined as adjusted p-value <0.05 with a Log2 fold-change cutoff of >0.4 for proteins and >1.0 for RNA. Significance determined by the Wilcoxon Rank Sum test with Bonferroni correction.

Analysis of both combined and individually annotated cell types showed significantly higher surface CD39 expression in children with SM compared to community children (**Fig. 6B-C**). Interestingly, upregulation of CD39 protein in children with SM appears to be post-transcriptionally regulated, as mRNA levels of *ENTPD1*, which is the gene encoding CD39^37^, remained unchanged across all cell types despite the surge in surface protein (**Fig. 6B-C**). This CD39 upregulation in children with SM was accompanied by a significant downregulation of ADA2 (*CECR1*) transcripts in plasmablasts (log2FC -1.3, p=0.025), the enzyme responsible for degrading extracellular adenosine^34^, compared to CC (**Fig. 6D**). The increased adenosine production via CD39 and decreased degradation via reduced ADA2 (*CECR1*) transcripts suggests a mechanism for adenosine accumulation in the clinical setting.

The T cell and myeloid compartments in children with SM also showed marked immunological dysregulation. The activated CD4^+^ T cell population (defined by CD223/LAG3^+^CD45RO^+^) displayed a transcriptional profile consistent with type 1 regulatory (Tr1) or exhausted T cells, characterized by the strong upregulation of LAG3, HAVCR2 (TIM-3), and CTLA4 (**Fig. 6E**). This lymphoid suppression was paralleled by a sepsis-like “paralysis” in the myeloid compartment, where monocytes downregulated MHC Class II genes (HLA-DQA2, log2FC -8.3) while upregulating the inflammatory alarmin S100A8 (log2FC 3.0) in children with SM (**Fig. 6F**)^38–40^. To confirm that these phenotypic shifts represented distinct, global cellular states rather than isolated marker changes, we generated unbiased heatmaps of the top 20 differentially expressed proteins and transcripts for each population (**Suppl. Fig. 6B-D**). The complete list of differentially expressed genes and proteins for all annotated cell types is provided in **Suppl. Table 1**. Together, these data support the presence of a systemic, purinergic-mediated immunosuppressive environment in children with severe malaria.

### Blockade of A2AR signaling ameliorates pathogenesis and restores humoral immunity

Our findings suggested that the gut microbiota might contribute to the immunosuppressive environment not only by inducing regulatory cells but also by providing a source of purine metabolites that leak into circulation upon barrier disruption. To test this, we performed targeted metabolomics on serum and cecal contents from mice at 0 and 7 dpi. Susceptible mice exhibited significantly higher basal and infection-induced levels of adenosine in both serum and cecal contents compared to resistant mice (**Fig. 7A**). While serum adenine levels increased in both groups upon infection, susceptible mice maintained a higher concentration in the cecum at 7 dpi (**Fig. 7B**). In contrast, resistant mice displayed significantly higher levels of the downstream catabolite uric acid, a known pro-inflammatory metabolite, in their serum both before and after infection (**Fig. 7C**). The metabolic intermediates inosine, hypoxanthine, and xanthine were largely depleted in the serum by 7 dpi (**Suppl. Fig. 7A**-**C**), suggesting rapid turnover of the pathway during the acute inflammatory response.

**Figure 7.**
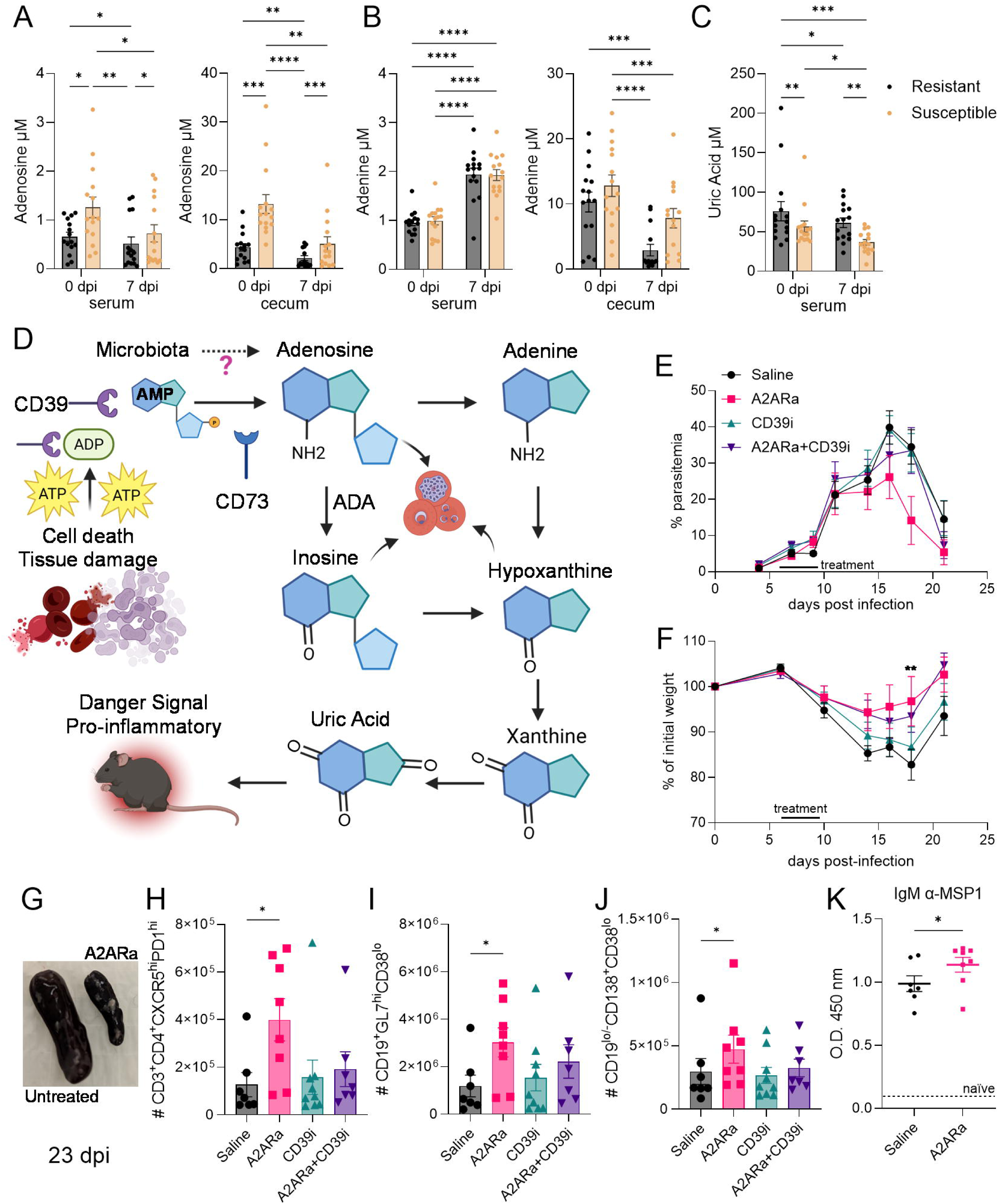
Purine metabolism drives pathogenesis, and A2AR blockade restores protective immunity. (**A**-**C**) Targeted metabolomics (LC-MS) of serum and cecal contents from Resistant and Susceptible mice (0 and 7 dpi), quantifying (**A**) Adenosine, (**B**) Adenine, and (**C**) Uric Acid. (**D**) Schematic of the purinergic signaling pathway, potential microbiota contributions, and downstream effects. Created with BioRender.com. (**E**-**K**) 8-12-week-old C57BL/6N mice were infected with *P. yoelii* and treated i.p. with an A2AR antagonist (8-3-chlorostyryl-caffeine 1 mg/kg), CD39i (ARL 67156 2 mg/kg) or both on days 6-9 post-infection. (**E**) Parasitemia curves were followed longitudinally. (**F**) Body weight was monitored longitudinally and compared to the initial weight. (**G**) Spleen image representation at 23 dpi. Mice spleens were collected for flow cytometry, gated on singlet live cells, and further gated as depicted in the Y axis for Tfh CD4 T cells (**H**), GC B cells (**I**), and antibody-producing cells (**J**). 23 dpi sera from mice were incubated with plates absorbed with *P. yoelii* Merozoite Surface Protein-1 (MSP-1) and tested with anti-IgM antibodies (**K**) and developed using our in-house ELISA. Statistics – Two-way ANOVA (A-C); Mann-Whitney (F-K). Median ± SEM. Data are representative of two independent experiments. *p<0.05, **p<0.01, ***p<0.001, ****p<0.0001.

These data support a model in which susceptible hosts exhibit higher systemic and cecal adenosine at early infection, potentially derived from the gut and exacerbated by the “emergency” plasmablast CD39+ response observed in sepsis-like-conditions (**Fig. 7D**)^29,41^. To functionally test this axis, we treated susceptible mice with an A2A receptor antagonist (A2ARa), a CD39 inhibitor (CD39i), or a combination of both during the peak of intestinal damage (days 6-9 p.i.). While treatment did not significantly alter the overall trajectory of parasitemia (**Fig. 7E**, **Suppl. Fig. 7D**) or survival (**Suppl. Fig. 7E**), blocking the A2A receptor significantly reduced disease severity. Mice treated with A2ARa exhibited improved sickness scores (**Suppl. Fig. 7F**-**G**) and were protected from severe weight loss (**Fig. 7F**, **Suppl. Fig. 7H**). This clinical improvement was associated with a marked reduction in splenomegaly at 23 dpi (**Fig. 7G**, **Suppl. Fig. 7I**), despite maintaining comparable total splenic leukocyte and T cell numbers (**Suppl. Fig. 7J**-**K**).

Critically, A2ARa-treated mice harbored significantly higher numbers of B cells (**Suppl. Fig. 7L**), Tfh cells (**Fig. 7H**), GC B cells (**Fig. 7I**), and plasma cells (**Fig. 7J**) compared to untreated controls at 23 dpi. This cellular recovery translated into a functional boost in humoral immunity, as A2ARa-treated mice produced significantly higher titers of anti-MSP1 and anti-*Py* extract IgM, but not IgG antibodies (**Fig. 7K**; **Suppl. Fig. 7M-N**).

Notably, inhibition of the upstream enzyme CD39, or a combination of both, failed to replicate these clinical or immunological benefits, suggesting that breakdown of extracellular ATP is essential for reducing pathology. Collectively, these results demonstrate that blocking A2AR signaling uncouples immunopathology from parasite burden and improves GC/Tfh architecture at 23 dpi, consistent with A2A involvement.

## Discussion

This study dissected the complex interplay between the gut microbiota, intestinal barrier integrity, and systemic immunity during *P. yoelii* infection, and disease severity. We demonstrate that a pre-existing gut microbiota-driven immune tone, characterized by elevated regulatory T cells, predisposes to hyperparasitemia. During infection, parasite-induced intestinal damage allows for detrimental gut-spleen crosstalk, culminating in an immunosuppressive, purinergic-driven emergency B cell response. The purinergic response is higher in children with severe malaria when compared to asymptomatic children. In mice, we show this cascade can be modulated by either reinforcing the gut barrier or blocking the downstream A2A receptor signaling, uncoupling severe pathogenesis from parasite burden.

Our work builds upon previous findings from our lab and others that established a link between gut microbiota composition and malaria outcomes in both mice and humans^4,9–12,42^. We previously showed that this susceptibility was associated with a contraction of the GC response by 9 dpi^16^. Here, we pinpoint an earlier checkpoint, demonstrating that susceptible mice fail to contract their splenic and gut-associated Treg populations at 7 dpi. This elevated regulatory environment is coupled with a systemically blunted IFN-γ, IL-10, and IL-4 response, creating an environment less conducive to the robust Tfh and GC B cell activation required for potent antibody production^21^. The expansion of regulatory T cells is known to contribute to malaria-induced immunosuppression, with studies showing that these cells impede protective GC responses through inhibitory molecules such as CTLA-4^43,44^. This landscape is further influenced by other IL-10-producing populations, such as Type 1 regulatory T (Tr1) cells, which also upregulate inhibitory receptors, like CTLA-4, during malaria but are associated with decreased disease severity in human cohorts^44^. This highlights a complex balance between immunosuppressive cell subsets, in which the regulatory environment that predisposes naïve hosts to severe acute malaria may, in the context of repeated exposures, evolve to mitigate immunopathology.

A central and surprising finding of our study is that while the basal gut immune environment confers opposing outcomes in a chemical colitis model (DSS), *Plasmodium* infection induces severe intestinal pathology and tight junction disruption in resistant and susceptible mice. This suggests that *P. yoelii* infection is a potent, independent driver of gut damage. It is an established clinical observation that severe malaria is a major risk factor for concurrent bacteremia caused by translocated enteric pathogens^7,11,42,45–56^. Our results reframe the role of the microbiota in this context: rather than causing the damage, specific microbial communities appear poised to exploit this infection-induced breach in the intestinal barrier. This relationship is bidirectional, as blood-stage malaria can induce a lasting dysbiosis that impairs host defense against malaria and secondary infections in mice^5,51,52^. This provides a mechanistic explanation for why simply colonizing germ-free mice with human microbiota from individuals who subsequently develop symptomatic malaria during the following malaria season can transfer the hyperparasitemia phenotype^14,15^. A recent study by Kroon et al. provides further mechanistic insight, showing that systemic LPS exposure in mice is sufficient to promote blooms of gut-luminal opportunistic pathogens by increasing gut-luminal oxygen species, which inhibit microbiota fermentation and fuel pathogen growth via oxidative respiration^57^. This observation fits within our recent findings demonstrating a higher abundance of pathogenic bacteria capable of oxidative respiration within the stool of children with severe malaria^13^.

Rather than being driven by pathobionts or a strictly species-specific phenotype, it appears that the immunomodulatory function of the microbiota is sufficient to recapitulate this phenotype, as we observed in multiple room barriers with different compositions over the years^12^. Through fecal microbiota transplantation, we functionally validated this concept. We demonstrated that the IgA-coated fraction of the microbiota from susceptible mice, composed not of classic pathogens but of highly immunomodulatory commensals and pathobionts^22^, was sufficient to induce the hyperparasitemia phenotype and suppress the splenic GC response in otherwise resistant hosts. Our observation of elevated *B. thetaiotaomicron*-tetramer+ T cells exclusively in gut-associated lymphoid tissues, rather than the spleen, points to an indirect mechanism of immunosuppression. This concept of gut inflammation breaking immune compartmentalization to alter responses at a distant site is supported by findings in other systems, where chemically-induced colitis was shown to disrupt established immune tolerance to commensal bacteria in the skin by promoting T cell trafficking and altering the local Treg/Teff balance^58^. The ability of this typically managed microbial community to drive pathology when translocated systemically highlights the critical principle that the detrimental potential of the microbiota is context-dependent and can be unleashed by a loss of intestinal barrier integrity in malaria.

We showed that this gut-spleen axis can be targeted therapeutically in our model. We improved clinical scores and survival in susceptible mice by administering the zonulin antagonist larazotide acetate to prevent intestinal barrier breakdown during the critical infection window. This intervention restored the splenic Tfh and GC B cell populations, effectively uncoupling the gut-driven immunopathology from the infection. Persistent increased permeability due to intestinal damage or an inability to repair tight junctions after insult can result in local and systemic inflammation, sepsis, septicemia, and multi-organ dysfunction in clinical patients^59,60^. Recently, Yonker and colleagues used larazotide acetate in a single patient, demonstrating proof-of-concept reductions in spike protein antigenemia and improved clinical status ^61^. A subsequent study using larazotide acetate as adjuvant therapy for MIS-C demonstrated the drug’s safety and tolerance in four additional children with MIS-C, as well as improved resolution of gastrointestinal symptoms^62^. The second study showed faster spike antigenemia clearance than the current standard of care for MIS-C^62^. These parallel findings in both experimental malaria and MIS-C suggest that losing barrier integrity can be a causal step in driving systemic immunopathology in acute inflammatory syndromes.

To understand the systemic consequences of this pathology, we investigated the purinergic signaling pathway, a critical regulator of inflammation and resolution^30^. A central mechanism is the rapid conversion of extracellular ATP from damaged cells into immunosuppressive adenosine via ectonucleotidases^31,32^. Our finding that susceptible mice mount an emergency B cell response characterized by an expansion of plasmablasts expressing the ectonucleotidase CD39 resonates with observations in sepsis^29^ and the “metabolic sink” phenotype in *P. yoelii* infection^63^. A link between the gut microbiota and CD39 expression is well-established in other contexts^64^; commensals such as *Bacteroides fragilis* are known to induce immunosuppressive CD39+ Tregs that protect against neuroinflammation in models of multiple sclerosis (EAE)^65^. The upregulation of this pathway is also translationally relevant, as increased expression of purinergic pathway components has been documented in patients with *P. vivax* malaria, and CD39 is also upregulated on antigen-experienced memory CD4^+^ T cells in children clinically immune to *P. falciparum*^66,67^. This phenotype is mirrored in our human cohort, where children with SM exhibit an upregulation of CD39+ in plasmablasts, among other cell types. Notably, our CITE-seq analysis revealed that this upregulation is post-transcriptional, as *ENTPD1* mRNA levels remained stable. Notably, while our murine model showed a compensatory systemic upregulation of ADA protein to manage purine overload, human plasmablasts from SM children exhibit lower CECR1 (ADA2) transcript levels. This transcriptional suppression implies a defect in the adenosine-degrading machinery during severe disease, potentially preventing the clearance of immunosuppressive metabolites.

The human data further contextualize this purinergic suppression within a broader state of immune paralysis. We observed that the expanded CD4+ T cell population in severe malaria patients expresses high levels of *LAG3*, *HAVCR2*, and *CTLA4*, a signature characteristic of Tr1 cells or exhausted T cells^44,68^. This aligns with our murine observation of increased regulatory responses and suggests that the “Treg” expansion we observe by flow cytometry likely includes this Tr1-like population. Simultaneously, monocytes in these patients exhibited a classic sepsis-like profile, characterized by upregulation of calprotectin (*S100A8/9*) and loss of the antigen presentation machinery (*HLA-DQA2*)^38,40,69^. As described by Nascimento *et al*. the CD39^hi^ plasmablasts can directly inhibit macrophage activity^29^. This underscores that the pathology of severe malaria is not merely a failure of B cells, but involves a purinergic-driven shutdown of both innate and adaptive arms, which our murine data suggest is driven by the gut-spleen axis.

To define the molecular signals transmitting this gut-driven suppression, we investigated the purine metabolome. We found that susceptibility was associated with the persistence of elevated adenosine levels in both the cecum and serum, identifying the gut as a potential reservoir for this nucleoside. Conversely, resistant mice exhibited significantly higher levels of the downstream catabolite uric acid, suggesting that their survival is linked to the efficient catabolism of immunosuppressive adenosine into pro-inflammatory danger signals. The dysregulation of purine metabolism is increasingly recognized as a hallmark of severe malaria. A recent study by Bond *et al*. (2025) identified hyperuricemia as a driver of acute kidney injury and mortality in children with severe malaria, linked mechanistically to hemolysis and intestinal injury^70^. Our data suggest a complementary pathogenic mechanism centered on the failure to clear adenosine, since mice (unlike humans) can degrade uric acid, reducing its toxicity. The presence of uricase-producing pathobionts in the gut, as noted by Bond *et al*., further complicates this landscape in humans, suggesting that the host-microbiota interplay strictly regulates the balance between adenosine-mediated suppression and uric acid-mediated toxicity.

The purinergic environment is not shaped solely by the host. *Plasmodium* spp. lack *de novo* purine biosynthesis and relies on the host salvage pathway^71^. Indeed, our data suggest that higher baseline adenosine levels in susceptible animals are associated with earlier and higher peak parasitemia. To secure these resources, infected red blood cells (RBCs) release ATP and engage host erythrocyte purinergic receptors (P2X, P2Y), thereby facilitating parasite development and activating new permeability pathways^72^.

Furthermore, the parasite expresses its own putative G-protein-coupled receptors, potentially allowing it to sense its microenvironment directly^73^. This suggests a synergy in which a parasite-driven signaling cascade, originating from the lysis of infected RBCs, potentiates and sustains the host’s gut-influenced purinergic suppressive program.

In our model, the greater adenosine presence despite high systemic ADA levels appears to act via A2AR signaling. Pharmacological blockade of the adenosine receptor A2AR, but not the upstream enzyme CD39, was sufficient to ameliorate disease severity and improve the GC and Tfh cell numbers and the anti-*Plasmodium* IgM response in susceptible mice. This suggests that excess adenosine acts as a key effector molecule. In contrast, the lack of efficacy of the CD39 inhibitor suggests that sufficient adenosine is already present, making receptor blockade a more effective therapeutic strategy. Further supporting the involvement of the purinergic pathway via A2AR, genetic polymorphisms in the ADORA2A gene itself have been associated with susceptibility to clinical *P. falciparum* malaria in human cohorts^74^.

In conclusion, our study proposes a multi-step model for microbiota-mediated malaria severity. A ’susceptible’ microbiota is associated with a high basal Treg tone.

Subsequent malaria infection induces gut barrier dysfunction, the translocation of microbial products or metabolites, such as adenosine. This systemic exposure triggers a pathological, purinergic-driven emergency immune response characterized by CD39⁺ plasmablasts and Tregs, which suppresses the development of a protective GC response and drives severe disease. Critically, our findings highlight the gut barrier and A2AR signaling as potent therapeutic targets. This suggests a promising adjunct strategy for severe malaria: reinforcing intestinal integrity (*e.g*., with larazotide acetate) or blocking adenosine signaling could be deployed alongside fast-acting anti-parasitic drugs to mitigate the immunopathology that causes mortality, potentially improving survival outcomes independent of direct effects on parasite burden.

## Limitations of the Study

While our study defines a critical mechanism linking gut barrier dysfunction to splenic immune paralysis, it also has limitations to consider. First, our human validation cohort, though deeply phenotyped via CITE-seq, comprised a relatively small sample size (n=8 per group). Larger longitudinal cohorts will be necessary to stratify patients based on specific purinergic biomarkers. Second, obtaining splenic tissue from children with severe malaria is not feasible; thus, we relied on PBMCs as a proxy for the systemic immune landscape. While phenotypic parallels between our mouse splenic data and human circulating cells were strong, the specific microenvironmental interactions within the human germinal center can only be inferred. Mechanistically, while we identified a failure to downregulate Tregs as a key correlate of susceptibility, we did not perform functional depletion studies to definitively establish their causal role in suppressing the germinal center response. Finally, while we demonstrate that the IgA-coated fraction of the microbiota drives this phenotype, we have not yet isolated the specific bacterial species or consortia responsible for the hyper-production of adenosine in this context. It also remains to be determined whether these “purinogenic” pathobionts directly secrete the ATP substrate (as seen with other commensals^75^) or if they induce host tissue damage that releases ATP for subsequent hydrolysis by CD39+ B cells. Future studies will be required to identify these specific pathways for potential therapeutic targeting.

## Methods

### Mice and Malaria Infection

C57BL/6N were sourced from different barrier rooms at Charles River Laboratories and Taconic Biosciences. Additionally, germ-free mice were colonized with cecal content from susceptible or resistant donor mice from the barrier rooms. Mice were housed in specific-pathogen-free conditions at the Indiana University School of Medicine. All experiments were performed following the guidelines and approval of the Indiana University School of Medicine Institutional Animal Care and Use Committee (IACUC).

For infections, frozen stocks of *Plasmodium yoelii* 17XNL were passed once in donor mice; blood was collected at 6 days post-infection and used to intraperitoneally inject experimental mice with 1.5x10^5^ infected red blood cells (iRBCs). Parasitemia was monitored by flow cytometry of peripheral blood stained with antibodies against Ter119 and CD45.2, along with Hoechst and Dihydroethidium (DHE) dyes^76^.

### Human Subjects and CITE-Seq Analysis

#### Study Population

Between 2014 and 2017, 600 children with SM were enrolled in the study designed to assess cognition in children < 5 years of age at 12 months follow-up, as described previously^35^. Children were eligible if they: (1) were between 6 months and 4 years of age; (2) had diagnostic evidence of malaria with either a positive rapid diagnostic test for *Plasmodium falciparum* histidine-rich protein-2 or direct visualization of parasites by Giemsa microscopy; (3) required hospitalization; and (4) had 1 or more of the following WHO SM criteria: coma (Blantyre coma score < 3), respiratory distress (deep acidotic breathing or lower chest wall retractions), multiple seizures (≥ 2 generalized seizures in 24 hours or a seizure > 30 minutes in duration), severe anemia (hemoglobin < 5 g/dL), or prostration (≥ 1 year, unable to sit unsupported or stand; < 1 year, unable to breastfeed). Children were recruited at 2 referral hospitals in Central and Eastern Uganda: Mulago National Referral Hospital in Kampala and Jinja Regional Referral Hospital in Jinja. Exclusion criteria included the following: known chronic illness requiring medical care, history of coma, head trauma, known developmental delay, cerebral palsy, or prior hospitalization for malnutrition. Delayed exclusion criteria included an elevated cerebrospinal fluid white blood cell count in a child with coma.

Community children, 120 total, were enrolled as controls recruited from the nuclear family, extended relatives, and nearby households, and were commonly enrolled during the one-month follow-up appointments for the children who had SM. Exclusion criteria included no illness requiring medical care within the previous four weeks and no findings of neurological abnormalities on physical exam, in addition to the other exclusion criteria previously stated. Malaria was evaluated in CC by either rapid diagnostic test for *Plasmodium falciparum* histidine-rich protein-2 or direct visualization of parasites by Giemsa microscopy followed by nested PCR targeting the 18S rRNA gene of *Plasmodium* spp., followed by amplification of *Plasmodium falciparum* using species-specific primers and visualization of amplicons though agarose gel electrophoresis, as previously described^77^, from genomic DNA isolation from whole blood using the QIAamp DNA Blood Mini Kit (Qiagen), according to the manufacturer’s instructions.

#### Ethics Approval and Consent to Participate

Initial verbal consent from the parents or legal guardians of study participants was obtained for children fulfilling inclusion criteria because most participants were critically ill and required emergency stabilization. Verbal consent was specified in the study protocol and approved by the respective IRB. Written informed consent was obtained once the participant was clinically stabilized. If a participant died before written informed consent could be obtained, verbal consent was considered sufficient. All study procedures and the sample collection schedule were outlined in the consent forms, and consent was obtained to store the samples. Ethical approval was granted by the institutional review boards at Makerere University School of Medicine, the University of Minnesota, and Indiana University. Regulatory approval was granted by the Uganda National Council for Science and Technology.

#### PBMC processing and CITE-Seq

Peripheral blood mononuclear cells (PBMCs) were isolated from whole blood by Ficoll-Paque density gradient centrifugation and cryopreserved in FBS with 10% DMSO. Samples were shipped from Uganda to the USA in a liquid nitrogen dry shipper.

Selection of the samples sent for single-cell analysis excluded children with HIV infection, positive parasites in stool, antibiotics prior to enrollment, and age < 2 years old. Samples were then randomly chosen to have equal representation of each sex within each study site (Kampala and Jinja) for children with severe malaria and community children (4 asymptomatic and 4 Pf. negative). Among the 8 children with severe malaria, 4 had cerebral malaria, and 4 had severe malaria anemia.

At Indiana University School of Medicine, the vials of PBMCs were thawed rapidly, washed in cold media, and assessed for cell viability. Dead cells were removed using a magnetic bead-based Dead Cell Removal Kit (Miltenyi Biotec). Live cells were then stained with a panel of oligo-conjugated antibodies (BioLegend TotalSeq-B Universal cocktail) and barcoded for CITE-seq analysis. Stained and labeled samples were pooled into four groups and run on High Throughput 10X Chromium with two replicates per pool by the Indiana University Medical Genomics Core. Raw sequencing data were processed using Cell Ranger to generate feature-barcode matrices. Downstream analysis was performed in R using the Seurat package (v5.2) and workflow. After quality control filtering and barcode-based demultiplexing, data from all samples were downsampled and combined into a single Seurat object. UMAP was used for dimensionality reduction and visualization. Cell type annotation was performed automatically using the Azimuth function with the pbmcref reference dataset of human PBMCs. Gene expression analysis was performed using the FindMarkers function on selected cell types, with transcript counts assessed using the Wilcoxon Rank-Sum test with Bonferroni correction. Plasmablast, monocyte, and activated T cell analyses were performed in children with severe malaria and community children, selecting on the predicted cell type; activated T cells were additionally selected for positive protein expression of CD223 (LAG3) and CD45RO, followed by gene expression analysis.

### In Vivo Treatments

FTY720 (Fingolimod): Mice were treated with 1 mg/kg FTY720 (Cayman Chemical) or vehicle control via intraperitoneal (i.p.) injection every other day from day -1 to day 14 post-infection.

DSS-induced Colitis: Mice were given 2% (w/v) dextran sulfate sodium (DSS, MW: 36,000-50,000, MP Biomedicals, Cat# 160110) in their drinking water for 7 days. Body weight was monitored on the depicted days.

Larazotide Acetate (L.A.): Mice were treated with 0.15 mg/mL L.A. in their drinking water and/or by oral gavage from days 6-10 post-infection.

A2ARa and CD39i Treatment: Mice were treated i.p. with 8-(3-chlorostyryl)-caffeine (A2ARa, 1 mg/kg), ARL 67156 (CD39i, 2 mg/kg), a combination of both, or saline vehicle from days 6-9 post-infection.

### Tissue Processing and Cell Isolation

Spleens, Peyer’s Patches, and mesenteric lymph nodes (MLNs) were harvested and processed into single-cell suspensions by mechanical disruption through a 70-µm cell strainer (Fisher Scientific, Cat: 22-363-548). Red blood cells in spleen samples were lysed using ammonium chloride potassium lysis buffer. Splenocyte suspensions were adjusted to a final volume of 3 mL in complete RPMI (RPMI 1640, 10% FBS, 1.19mg/mL HEPES, 0.2 mg/mL L-glutamine, 0.05 mM 2-mercaptoethanol), and 25 µL aliquots were used for each staining panel with equal volume acquired in the flow cytometer Attune NxT (Thermo Fisher Scientific). For MLNs and Peyer’s Patches, the entire single-cell suspension was divided equally among the required staining panels.

### Flow Cytometry

Before surface staining, single-cell suspensions were stained with a viability dye Zombie Aqua (BioLegend, Cat: 423102). For intracellular staining of Foxp3 and cytokines, cells were stained with the depicted extracellular markers, fixed, and permeabilized using Foxp3/Transcription Factor Staining (eBioscience, Cat: 00-5523-00) and Cyto-Fast^TM^ (Biolegend, Cat: 426803), respectively. For all samples, equal acquisition volumes were collected on an Attune NxT Flow Cytometer (Thermo Fisher), and total cell numbers were back-calculated (spleen diluted 1:100, resuspended in FACS buffer, and 85% of the total sample volume acquired). Data were analyzed using FlowJo^TM^ software (v10, BD Life Sciences). For unsupervised analysis, 10,000 live CD19+ events per mouse (5 per group) were concatenated and analyzed using the FlowSOM^33^ and t-SNE algorithms in FlowJo.

### Cytokine and Antibody Quantification

Serum for cytokine analysis (with protease inhibitor cocktail; Sigma-Aldrich, Cat: P8340) and antibody analysis was obtained via retro-orbital bleed on the indicated days, allowed to clot for 30-60 minutes, cleared by centrifugation, and stored at -80 °C until use.

Spleen Homogenates: Spleens were processed using the Bead Rupture Elite Homogenizer (Omni International) in ice-cold PBS with protease inhibitor cocktail (Sigma-Aldrich, Cat: P8340). Supernatants were collected after centrifugation and stored at -80°C.

LEGENDplex and ELISA: IFN-γ and IL-10 levels in spleen supernatants and serum were quantified using a LEGENDplex Mouse Inflammation Panel (BioLegend). IL-21 (ELISA MAX^TM^, Cat: 446107) and IL-4 (ELISA MAX^TM^, Cat: 431104) were measured by individual ELISAs (Biolegend). Serum levels of TFF3, FABP2, soluble CD14, LBP and NGAL were quantified by ELISA.

To evaluate MSP-1_19_-specific or iRBC lysate-specific antibodies, MaxiSorp Immuno plates (Thermo Fisher Scientific, Waltham, MA) were coated with 0.5 µg/ml recombinant MSP-1_19_ or 10 µg/mL iRBC lysate overnight at 4°C. Plates were blocked for 2 hours at room temperature (RT) with 2.5% w/v BSA + 5% v/v FCS in PBS. Dilutions of serum were added to wells and incubated 3 hours at room temperature. Horseradish peroxidase-conjugated goat anti-mouse IgM or IgG (Jackson ImmunoResearch, West Grove, PA) was added and incubated for 1 hour at RT. Plates were developed with a TMB substrate set (Biolegend, San Diego, CA). Two molar H_2_SO_4_ was used to halt the reaction, and plates were read using a microplate reader with the absorbance read at an absorbance 450 nm.

### Microbiota Analysis

IgA-SEQ: The IgA-SEQ protocol was adapted from Palm *et al*. (2014)^22^. Cecal contents were incubated with PE-conjugated anti-mouse IgA, followed by anti-PE MicroBeads Ultrapure (Miltenyi Biotec, Cat: 130-105-639) for enrichment. DNA was extracted from pooled fecal pellets prior to (input), IgA-positive (IgA), and flow-through (FT) fractions using the QIAamp PowerFecal Pro DNA Kit (Qiagen). DNA concentration was measured using Qubit 3 Fluorometer (Invitrogen) and sent to the Indiana University Center for Medical Genomics for whole-genome shotgun metagenomics sequencing and library preparation on NovaSeq 6000 with 150 bp paired-end sequencing. Quality control, host sequence removal, and genome assembly were completed with the mNGS Illumina pipeline by CZ ID78 to produce a feature table with species-level bacteria for each sample. The feature table was imported into R and converted to a phyloseq object.

Bar plots representing the top 10 bacteria at the family and species levels for each group were merged across the three days and visualized using the MicroViz package^78^.

Fecal Microbiota Transplant (FMT): IgA-enriched or flow-through fractions from susceptible or resistant donors were gavaged into resistant recipient mice for three consecutive days, one week before infection. For cytometric validation of purity, stained bacteria were suspended in 100 µL of ice-cold saline (0.9% NaCl, 0.1 M HEPES, pH 7.0) containing 5 µM SytoBC (Invitrogen, Cat: S34855) and incubated for 15 minutes on ice in the dark. DAPI (0.5 µg/mL) was added to distinguish live from dead bacteria.

Samples were diluted with 500 µL staining buffer, filtered through a 70 µm filter cap, and acquired on an Attune NxT Flow Cytometer (Thermo Fisher). Bacterial events were gated on SytoBC+ and DAPI-populations and gavaged above 75% IgA+.

### Histology and Immunofluorescence

Small intestines were rolled, mounted in cassettes, and fixed in 10% neutral buffered formalin and embedded in paraffin. 7-µm sections were mounted on glass slides and H&E-stained. For the semi-quantitative analysis of small intestinal damage, the slides were digitized using Aperio Whole Slide Digital Imaging and visualized with Aperio ImageScope Software (Leica Biosystems). Low and high-magnification objectives were used. The score for analyzing inflammation and intestinal alterations is available in **Table 1**.

**Table 1.** Histopathological scoring system for mouse intestinal inflammation.

|  |  | Histopathological findings |
| --- | --- | --- |
| Score |  | Observations |
| 0 | Absent | No histopathological changes. |
| 1 | Slight | Slight edema in the mucosa and/or dilation of the galactophorous duct, with few inflammatory cells. |
| 2 | Moderate | Mucosa exhibiting mild to moderate inflammatory infiltrate, with damage to the enterocytes. |
| 3 | Marked | Mucosa presenting moderate inflammatory infiltrate with damage to the enterocytes or focal erosions, and the presence of mild to moderate infiltrate in the submucosa. |
| 4 | Severe | Mucosa presenting intense inflammatory infiltrate, with damage to the enterocytes or focal erosions, and moderate to intense infiltrate in the submucosa. |

For immunofluorescence, 0.5-cm sections the distal part of the small intestines was embedded in OCT, cryosectioned (7-µm), mounted in glass slides, frozen, methanol fixed and stained with antibodies against ZO-1 Polyclonal Antibody (ZMD.437, Thermo Fischer, Cat: 40-2300) and Occludin (OC-3F10, Thermo Fischer, Cat: 33-1500) and mounted using DAPI (Vectashield® with DAPI, Cat: H-1200-10). Images were acquired on a Zeiss LSM800 Microscope and analyzed using Zen Blue Lite 3.12.

### Intestinal Permeability Assay

Mice were fasted for 4 hours and then gavaged with 4 kDa FITC-dextran (Sigma-Aldrich, Cat: 46944). Blood was collected 4 hours later. FITC fluorescence in the serum was quantified using a spectrophotometer (SpectraMax M2) and compared to a standard curve of FITC-dextran diluted in normal mouse serum.

### Targeted Metabolomics (UPLC-MS/MS)

To prevent rapid enzymatic degradation of adenine nucleotides during sample collection, blood was collected in tubes containing 10 mM pentostatin (an adenosine deaminase inhibitor, Tocris). Samples were centrifuged at 1,000 x *g* for 15 minutes at 4°C, and the resulting plasma was stored at -80°C until extraction. Cecal contents were harvested by flushing the cecum with PBS containing pentostatin and stored at -80°C.

Protein removal and metabolite extraction were performed using a methanol precipitation protocol adapted for UPLC-MS analysis. Briefly, 100 µL of plasma or cecal suspension was combined with 300 µL of ice-cold HPLC-grade methanol (Fisher Scientific). Samples were vortexed for 30 seconds and incubated at -20°C for at least 30 minutes to facilitate protein precipitation. Following incubation, samples were centrifuged at 14,800 rpm for 10 minutes at 4°C. A 360 µL aliquot of the supernatant was transferred to a fresh centrifuge tube, discarding the pellet. The supernatant was dried completely using a Vacufuge Plus (Eppendorf) for approximately 2 hours. Dried samples were reconstituted in 90 µL of pure water. The concentrations of adenosine, adenine, inosine, hypoxanthine, xanthine, and uric acid were quantified by Ultra-Performance Liquid Chromatography-Tandem Mass Spectrometry (UPLC-MS/MS).

### Serum Protein and Adenosine Deaminase Quantification

Serum adenosine deaminase (ADA) levels were measured using a mouse Adenosine Deaminase ELISA kit (LSBio, Cat: LS-F32097-1). Serum levels of lipopolysaccharide-binding protein (LBP) (Hycult Biotech, Cat: HK20502), soluble CD14 (R&D Systems, Cat: DY982), angiopoietin-2 (Fisher Scientific, Cat: MANG20), and ICAM-1 (Fisher Scientific, Cat: EMICAM1ALPH) were quantified according to the manufacturers’ instructions.

### Statistical Analysis

Statistical analyses were performed using GraphPad Prism (v10). Specific tests include the Mann-Whitney U test, one-way ANOVA with Tukey’s multiple comparison test, Two-Way ANOVA for longitudinal comparisons, and the Mantel-Cox Log-Rank test for survival curves. Data are presented as mean ± SEM. P-values < 0.05 were considered significant.

## Data and Materials Availability

The datasets generated and analyzed during the current study are available in the NCBI Sequence Read Archive, pending dbGaP/PHS/IRB; WGS bacteria: pending. All data will be made available upon request.

## Codes Availability

Codes are available on https://github.com/pending.

## Declaration of Generative AI and AI-Assisted Technologies in the Writing Process

During the preparation of this work, the authors used Gemini Advanced to improve the readability and flow of the manuscript and to review the text for potential overstatements or lack of clarity. Pre-submission review was conducted using qed Science (https://www.qedscience.com). Additionally, AI assistance was utilized to refine Python scripts for the specific customization of volcano plots and heatmaps (Figure 6, Suppl. Figure 6). The authors reviewed and edited all AI-generated content and take full responsibility for the accuracy and integrity of the final manuscript and data visualization.

## Acknowledgements

We thank Dr. Renan de Carvalho for helpful discussions on B cell populations and gating, Dr. Danielle Nascimento for her advice on the preparation and administration of adenosine-related drugs, and Morgan Little for technical assistance. We thank Drs. Prasida Holla and Jyoti Bhardwaj for assistance with the CITE-seq preparation. We thank Dr. Sabrina Absalon for her assistance and for providing access to the LSM900 microscope. We thank the NIH Tetramer Core Facility (NIH Contract 75N93020D00005 and RRID: SCR_026557) for providing SusC peptide and scramble peptide tetramers, and Dr. Scott Lindner for the recombinant anti-MSP-1 antibody. We thank the IUSM Flow Cytometry Cancer Center Core for their assistance. Metagenome sequencing of fecal bacteria and sequencing of CITE-seq samples, and CellRanger_Counts analysis were carried out in the Center for Medical Genomics at Indiana University School of Medicine, which is partially supported by the Indiana University Grand Challenges Precision Health Initiative. Histology services were provided by the Histology Core of the Indiana Center for Musculoskeletal Health at the IU School of Medicine and the Indiana Clinical and Translational Sciences Institute (CTSI).

## Author Contributions

**R.B.P.** conceptualized and designed the study, performed most experiments, analyzed data, created figures, and wrote the original draft. **O.J.B., F.M.S.O., E.F., S.L.,** and **A.T-L.** contributed to methodology, investigation, formal analysis, and visualization. **L.B., M.R.F.,** and **S.K.** contributed to experimental investigation and data curation. **R.N., R.O.O.,** and **C.J.** provided critical patient samples and associated clinical data. **J.B.** performed and interpreted metabolomics data. **J.C.A-F.** provided guidance and conceptual advice on the purinergic pathway. **N.W.S.** secured funding, supervised the project, contributed to conceptualization, and reviewed and edited the manuscript. All authors reviewed, provided feedback on, and approved the final manuscript.

