## Supplemental Figures for "Intestinal Barrier Loss Enables Microbiota-Mediated Purinergic Suppression During Malaria"

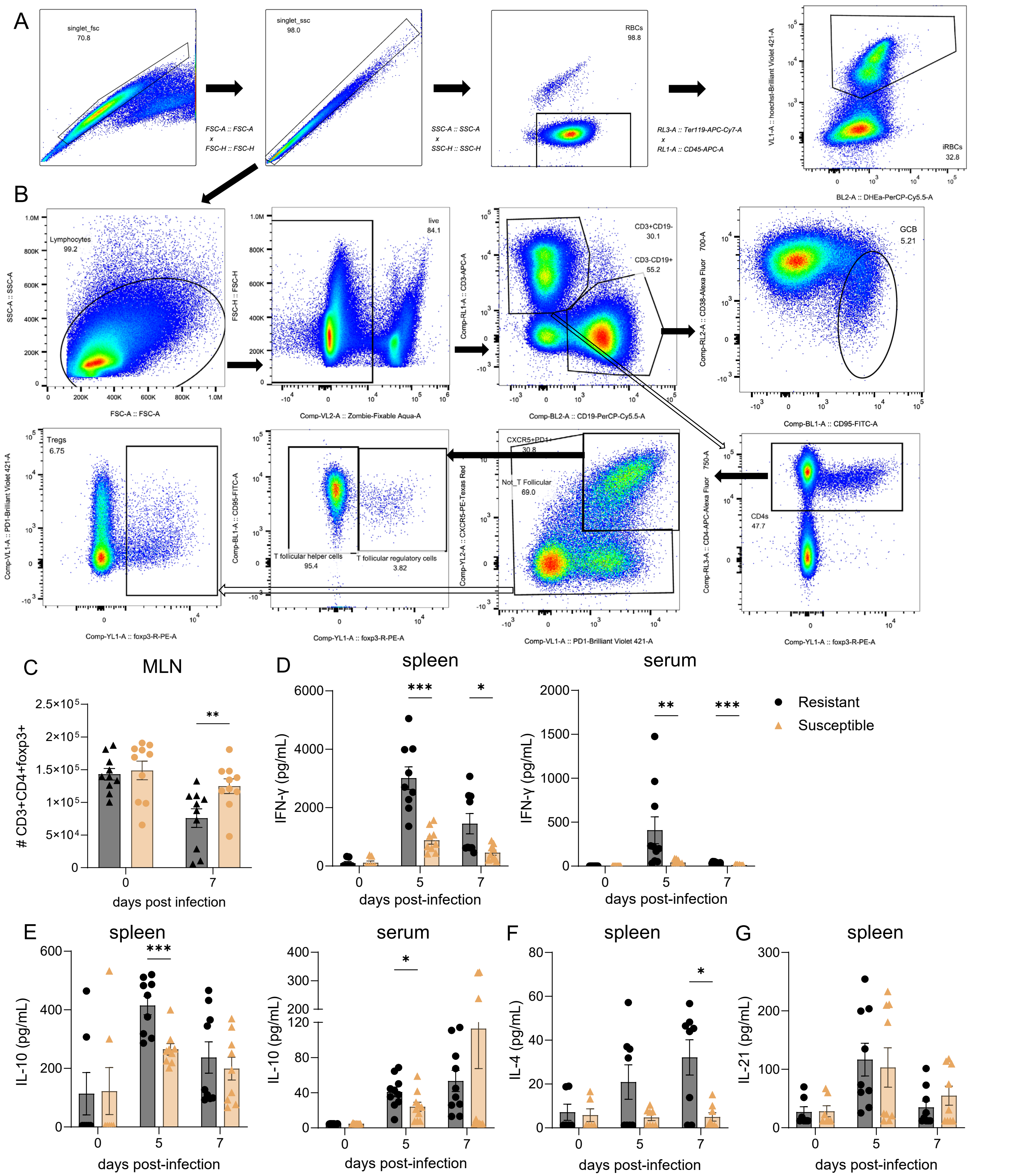


**Supplementary Figure 1. Susceptibility to hyperparasitemia is associated with blunted systemic cytokines.** (**A**) Representative flow cytometry gating strategy for quantifying parasitemia in peripheral blood using Hoechst and TER-119 staining. (**B**) Representative gating strategy for immunophenotyping of splenic T cells and B cells. (**C**) Mesenteric lymph nodes (MLNs) Resistant and Susceptible mice were removed before infection (0), and 7 days p.i., and processed for flow cytometry. Cells were gated as in (B) to identify Regulatory T cells. (**D-E**) Quantification of (**D**) IFN-γ and (**E**) IL-10 in spleen homogenates and serum by LEGENDplex. (**F-G**) Quantification of (**F**) IL-4 and (**G**) IL-21 in spleen

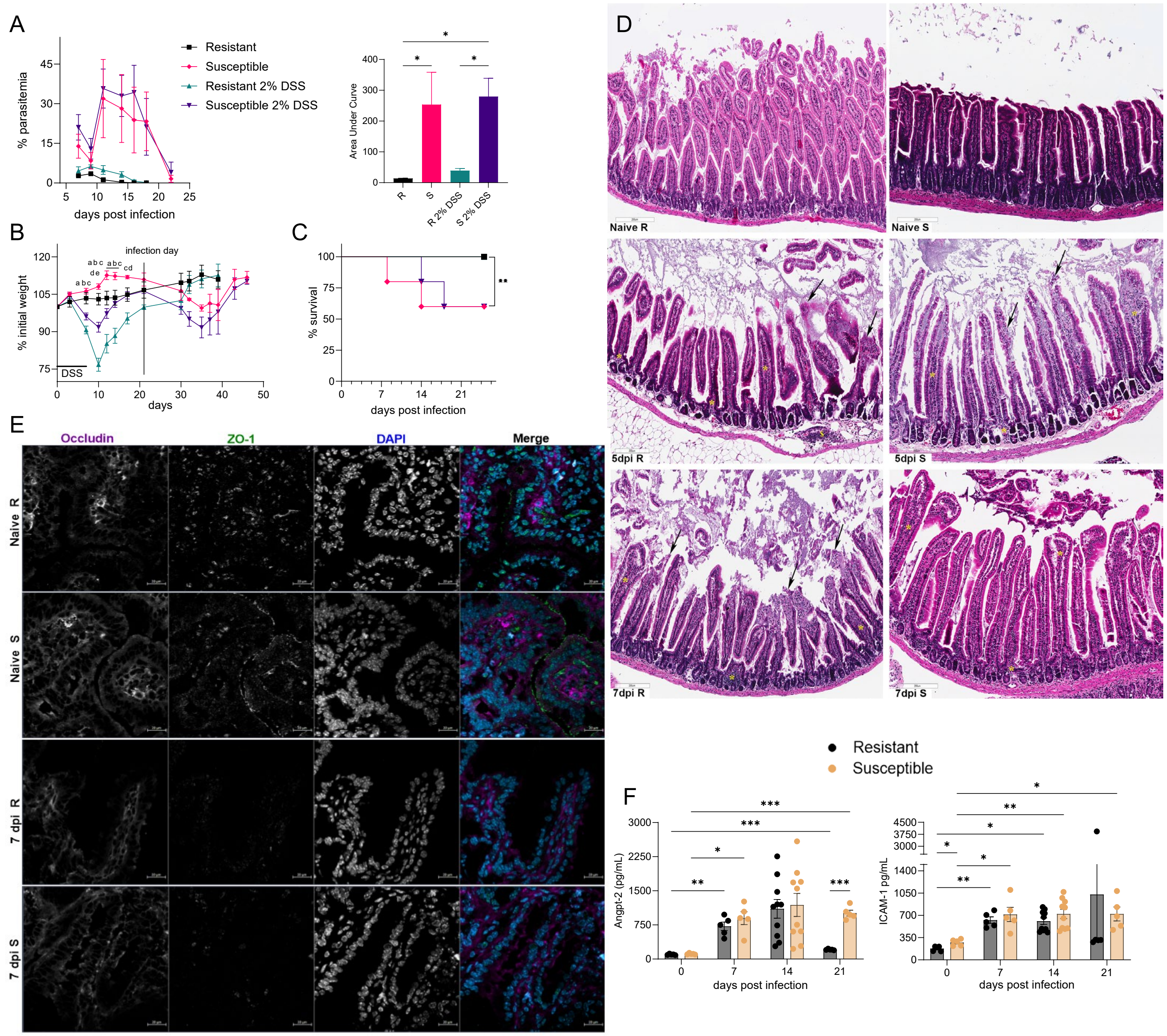


**Supplementary Figure 2. Transient DSS-induced dysbiosis does not alter malaria outcome, and infection induces vascular markers systemically.** Resistant and Susceptible mice were treated with 2% DSS for 7 days and allowed to recover their initial body weight before being infected with *P. yoelii*. (**A**-**C**) Parasitemia and AUC (**A**), percent of initial weight (**B**), and survival (**C**) were monitored. (**D**) H&E histology of small intestine at d0, d5, and d7. (**E**) Representative individual channel and merged immunofluorescence images of the distal small intestine stained for Occludin (purple), ZO-1 (green), and DAPI (blue). (**F**) Longitudinal serum quantification at the indicated timepoints and subjected to ELISA to quantify Angiopoietin-2 (Angpt-2) and ICAM-1. Statistics - Mann-Whitney tests. N=5, except 14 dpi (n=10), mean ± SEM (A).Results are representative of two or more experiments.

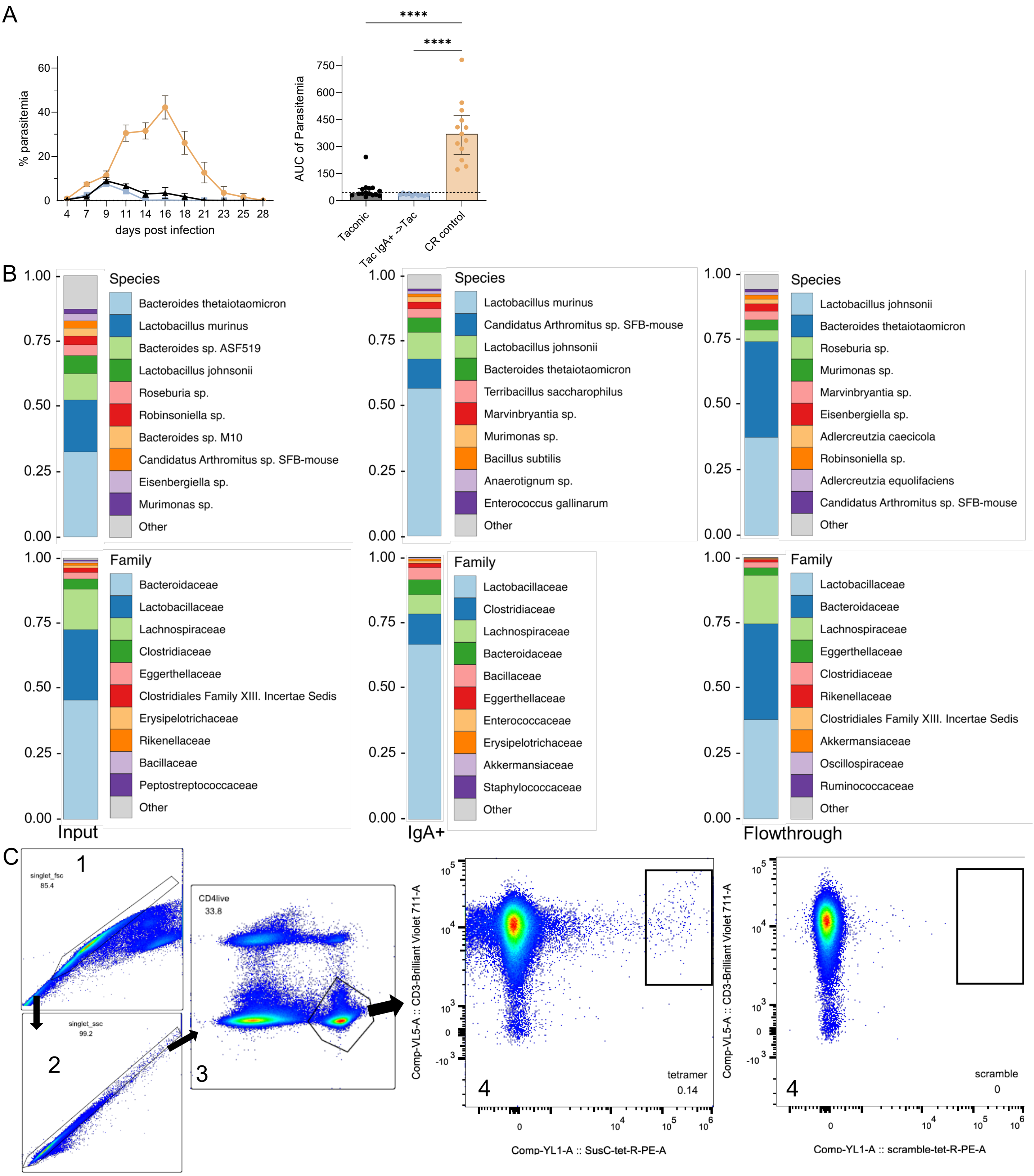


**Supplementary Figure 3. Characterization of microbiota fractions and control fecal transplant.** (**A**) Parasitemia and Area Under the Curve (AUC) of resistant mice gavaged with IgA-enriched fractions from **resistant** donors (control), showing no transfer of susceptibility. (**B**) Taxonomic composition (16S/Metagenomic, Species and Family) of the Input, IgA-enriched, and Flow-through fractions, confirming distinct microbial profiles and successful enrichment of IgA-coated taxa. (**C**) Representative flow cytometry gating strategy for *Bacteroides thetaiotaomicron* SusC-like tetramer-positive CD4+ T cells. Data are presented as mean ± SEM. Significance by One-Way ANOVA.

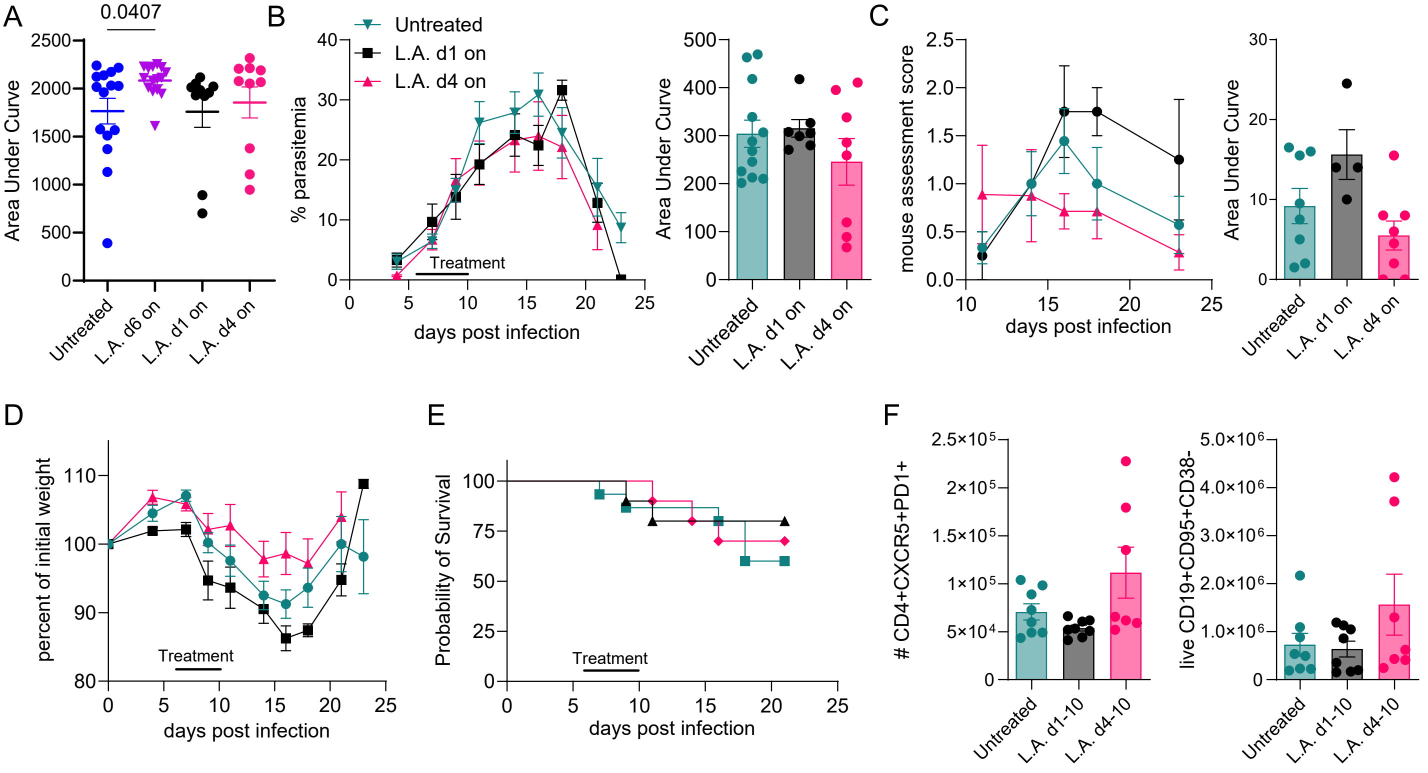


**Supplementary Figure 4. Therapeutic window of Larazotide Acetate treatment.** (**A**) Area under the curve (AUC) for the weight loss curves shown in Figure 5D. (**B-E**) Susceptible mice were infected with *P. yoelii* and treated with Larazotide Acetate from days 1-10 or 4-10 post-infection. Parasitemia (**B**), clinical scores (**C**), weight (**D**) and survival (**E**) were monitored. Twenty-three days post-infection, mice spleens were collected, and $2\times10^6$ single splenocytes were stained with Zombie Aqua (BioLegend) to distinguish live and dead cells, washed, and stained with the depicted antibodies for immunophenotyping on an Attune NxT Cytometer. (**F**) Cells were gated as single cells, then lymphocytes, live (Zombie Aqua negative) CD3+, and then gated as depicted for T Follicular CD4 T cells and Germinal Center B cells. Pairwise comparisons for AUC and cell frequencies were compared using the Mann-Whitney unpaired test (two-tailed). Initial N=15 for untreated and 10 for treated days 1-10 and 4-10 except for C (N=10 for untreated, d4-10 and 5 for d1-10).

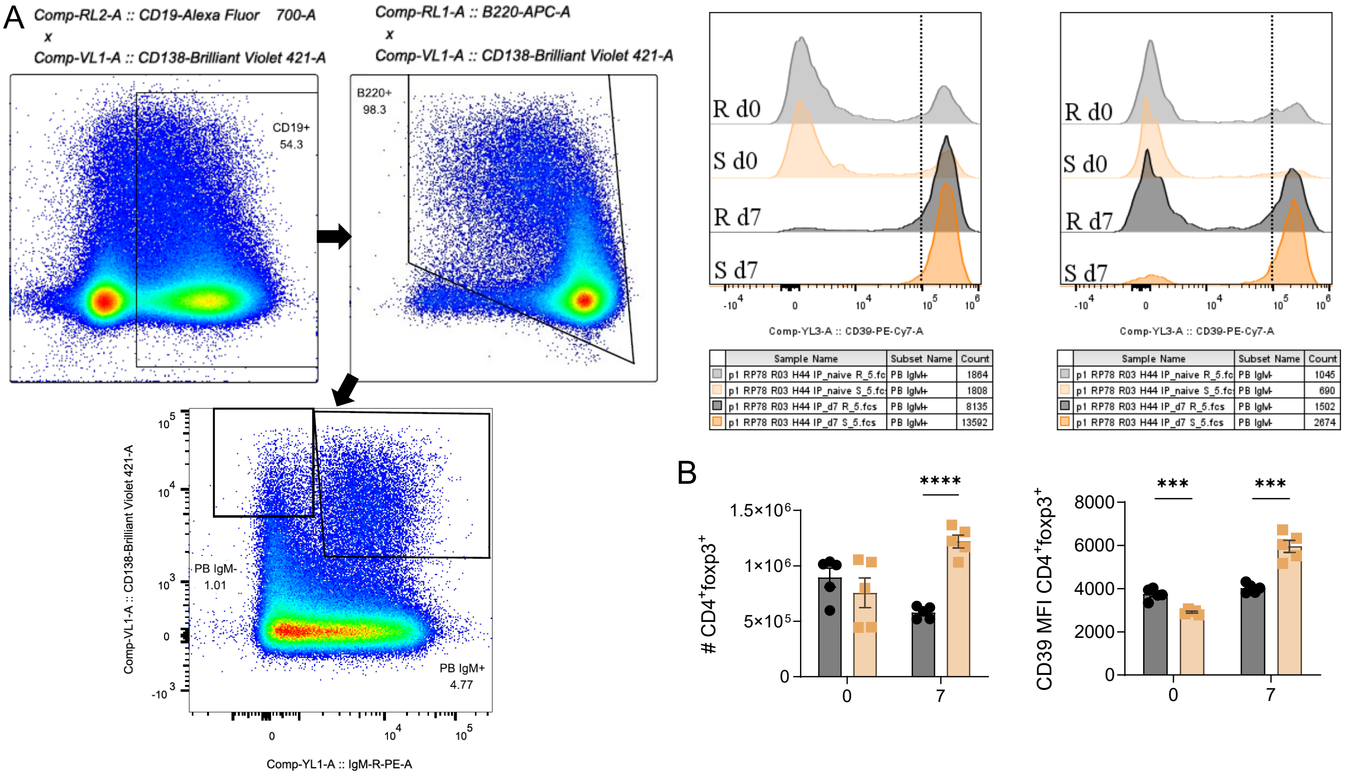


**Supplementary Figure 5. CD39 expression on CD138+ cells and Tregs. (A)** Gating strategy and representative histograms for CD39 MFI on IgM+ and IgM- plasmablasts after gating single live cells as in Suppl. Fig. 1A-B. (**B**) Quantification of splenic Treg numbers (as gated in Suppl. Fig. 1B) and CD39 MFI on Tregs before and after infection. Statistics - Mann-Whitney test. Data are representative of two independent experiments. ***p<0.001, ****p<0.0001.

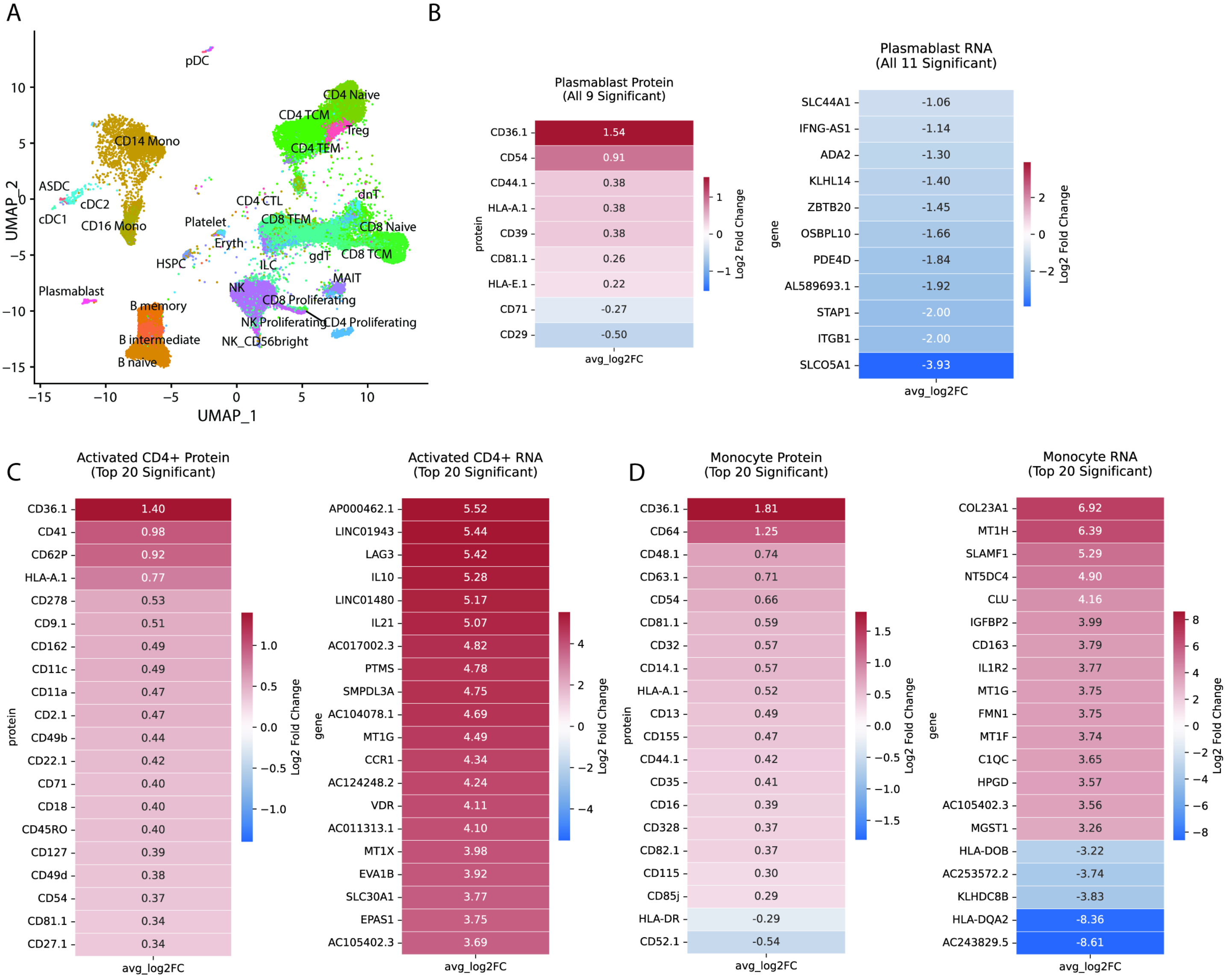


**Supplementary Figure 6. Transcriptional and proteomic signatures of immune exhaustion and myeloid paralysis in severe malaria.** (**A**) UMAP visualization of human PBMCs from all donors (n=8 Severe Malaria, n=8 Community Children), annotated by major cell type clusters. This map serves as the reference for the split UMAPs shown in Fig. 6A. (**B**-**D**) Heatmaps displaying the top 20 differentially expressed surface proteins (left columns) and mRNA transcripts (right columns) for (B) Plasmablasts, (C) Activated CD4+ T cells, and (D) Monocytes comparing Severe Malaria (SM) versus Community Controls (CC). Features are ranked by average log2 fold-change (log2FC), where red indicates upregulation and blue indicates downregulation in SM relative to CC. The color bar represents the scaled average log2FC. Significance determined by Wilcoxon Rank Sum test with Bonferroni correction.

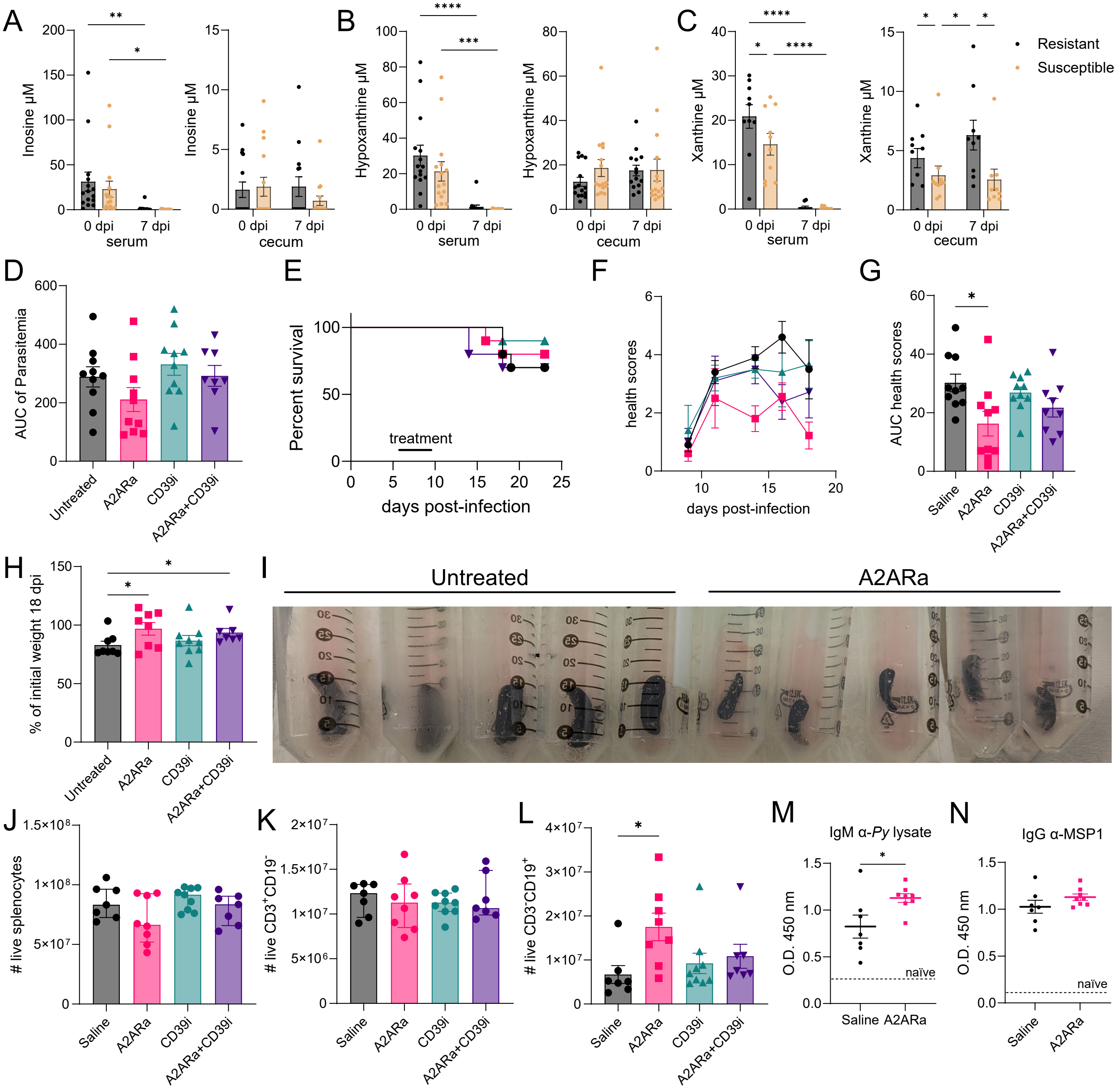


**Supplementary Figure 7. Purine metabolites and A2ARa treatment efficacy.** (**A-C**) LC-MS quantification of (**A**) Inosine, (**B**) Hypoxanthine, and (**C**) Xanthine in serum and cecum. (**D**) AUC of Parasitemia for treated groups. (**E**) Survival curves. (**F-G**) Health scores and AUC of health scores. (**H**) Weight at 18 dpi. (**I**) Representative spleen images. (**J-L**) Total counts of (**J**) Live splenocytes, (**K**) CD3+ T cells, and (**L**) CD19+ B cells. (**M**) Anti-*P. yoelii* lysate IgM. (**N**) Anti-MSP1 IgG levels. Statistics – Two-way ANOVA (A-C); Mann-Whitney (D-N). Median ± SEM. Data are representative of two independent experiments. *p<0.05, **p<0.01, ***p<0.001, ****p<0.0001.
